# De Novo Protein Design for Novel Folds using Guided Conditional Wasserstein Generative Adversarial Networks (gcWGAN)

**DOI:** 10.1101/769919

**Authors:** Mostafa Karimi, Shaowen Zhu, Yue Cao, Yang Shen

## Abstract

**Motivation:** Facing data quickly accumulating on protein sequence and structure, this study is addressing the following question: to what extent could current data alone reveal deep insights into the sequence-structure relationship, such that new sequences can be designed accordingly for novel structure folds?

**Results:** We have developed novel deep generative models, constructed low-dimensional and generalizable representation of fold space, exploited sequence data with and without paired structures, and developed ultra-fast fold predictor as an oracle providing feedback. The resulting semi-supervised gcWGAN is assessed with the oracle over 100 novel folds not in the training set and found to generate more yields and cover 3.6 times more target folds compared to a competing data-driven method (cVAE). Assessed with structure predictor over representative novel folds (including one not even part of basis folds), gcWGAN designs are found to have comparable or better fold accuracy yet much more sequence diversity and novelty than cVAE. gcWGAN explores uncharted sequence space to design proteins by learning from current sequence-structure data. The ultra fast data-driven model can be a powerful addition to principle-driven design methods through generating seed designs or tailoring sequence space.

**Availability:** Data and source codes will be available upon request.

**Supplementary information:** Supplementary data are available at *Bioinformatics* online.

## 1 Introduction

A fundamental science question about proteins, the workhorse molecule of life, is their sequence–structure–function relationships (Alberts *et al*., 2015). Anfinsen and co-workers studied the renaturation of fully denatured ribonuclease (Anfinsen *et al*., 1962), and eventually established a thermodynamic hypothesis (Anfinsen, 1973). Since then, the direct exploration of sequence–structure relationship has led to both the forward problem of structure prediction from sequence (Baker and Sali, 2001) as well as the inverse problem of sequence design for desired structures (Pabo, 1983; Street and Mayo, 1999). With the data quickly accumulatinng on protein sequence and structure, a central question this study is addressing is: to what extent could current data alone reveal deep insights into sequence-structure relationships to empower protein design?

The forward problem of protein structure prediction, especially *ab initio* prediction without templates, is often solved by energy minimization. Even this classical principle-driven approach has benefited from data. Examples include the use of structural fragments for efficient sampling and the use of structure and sequence data for training scoring functions. A recent wave of data comes from protein sequences without paired structures. Specifically, sequence coevolution can be exploited to infer residue-residue structure contacts (Weigt *et al*., 2009; Kamisetty *et al*., 2013; Seemayer *et al*., 2014) and enhance protein structure prediction significantly (Kim *et al*., 2014; Hopf *et al*., 2014; Ovchinnikov *et al*., 2014). As witnessed in recent CASP (Critical Assessment of Structure Prediction) rounds, the latest revolution is in the prediction of residue-residue distances even for proteins with few homolog sequences, which is enabled by advanced deep neural network architectures (especially deep residual networks) that learn from the sequence, structure, and co-evolution data (Wang *et al*., 2017)

The inverse problem of protein design is often similarly pursued following the energy minimum principle (Koga *et al*., 2012; Gainza *et al*., 2016; Pierce and Winfree, 2002). Current protein (re)design algorithms fall in three classes: 1) exact algorithms such as dead-end elimination, A*, and cost function networks (Gainza *et al*., 2013; Traoré *et al*., 2013; Hallen and Donald, 2016; Karimi and Shen, 2018); 2) approximation algorithms such as relaxed integer-programming and loopy belief propagation (Kingsford *et al*., 2004; Fromer and Yanover, 2008); and 3) heuristic algorithms such as genetic algorithms and Markov chain Monte Carlo (MCMC) (Jones, 1994; Leaver-Fay *et al*., 2011). Using multiple backbone “blueprints” from fragments or geometry, they could be extended to *de novo* protein design where even the exact backbone structure is assumed unknown along with the sequence (Huang *et al*., 2016). In particular, the (energy minimum) principle-driven Rosetta tools (Leaver-Fay *et al*., 2011) have made great success for *de novo* protein design (Marcos *et al*., 2017; Shen *et al*., 2018; Dou *et al*., 2018; Silva *et al*., 2019).

In contrast to the forward problem, the inverse problem of *de novo* protein design has witnessed limited impacts from deeply exploiting data with advanced artificial intelligence (especially deep learning) technologies (Greener *et al*., 2018; Ingraham *et al*., 2019). Meanwhile, the impacts of deep generative models, represented by Generative Adversarial Networks (GAN) (Goodfellow *et al*., 2014) and Variational Auto-Encoder (VAE) (Kingma and Welling, 2013), have reached the sibling fields of inverse design for DNA (Gupta and Zou, 2018; Killoran *et al*., 2017), RNA (Eastman *et al*., 2018), small molecules (Popova *et al*., 2018; Guimaraes *et al*., 2017), and peptides (Muller *et al*., 2018).

What are the unique challenges for deep generative models in *de novo* protein design for a novel fold, the topic of this study? They come from the design space, the design objective (desired properties), and the mapping in between. The first challenge, a numerical one, comes from the much more daunting protein sequence space. Compared to aforementioned molecular designs, protein sequences have more choices at each position (20 standard amino acids versus 4 nucleotides) and are much longer, leading to the dimensionality of 20^*L*^ ≫ 4^*K*^ where *L > K*. The second challenge, a conceptual and mathematical one, is that, the fold space is a discrete domain that has not been completely observed (Kolodny *et al*., 2013). Therefore, a generalizable representation is needed to design a novel fold (a value in the discrete space) never seen in training data. In contrast, aforementioned deep generative small-molecule designs often target either a continuous property (such as logP) or a discrete one with desired values observed in training data. The last challenge is the knowledge gap about the complex sequence-fold relationship. Protein folds are products of both convergent and divergent evolution (Gu and Bourne, 2009). Very similar sequences often have the same fold with not-so-rare exceptions; and very dissimilar sequences can have the same fold. In contrast, designing RNAs benefits from the fact that desired structures can often be readily translated to base pairing patterns in the sequence space.

We present a study aiming at exploiting current data and advanced computing technologies for faster, broader, and deeper exploration of the protein sequence space while seeking principles underlying protein structure folds. To overcome the aforementioned challenges in protein design, we have developed a semi-supervised, guided conditional Wasserstein GAN (gcWGAN) by exploiting 1) a generalizable low-dimensional fold representation, 2) protein sequence data without paired structures, and 3) feedback from a fold predictor (or “oracle”). Specifically, we first use kernel Principle Correlation Analysis (kPCA) on 1,232 known folds to learn fold representations generalizable for novel folds. We then develop a novel Generative Adversarial Network (GAN) for protein sequence generation. The model is conditioned on the low-dimensional fold representation and guided through the oracle. It is pre-trained using abundant unsupervised protein sequence data (without structures) and fine-tuned using limited supervised sequence data. We systematically assess our models’ capability on designing novel protein folds, including a newly published one. We find that, compared to a recent study based on conditional VAE (Greener *et al*., 2018), our models generate proteins that are comparably or more accurate in desired folds, yet much more diverse and often more novel in sequence.

The architecture of gcWGAN is illustrated in Fig. 1 and detailed next.

**Fig. 1:**
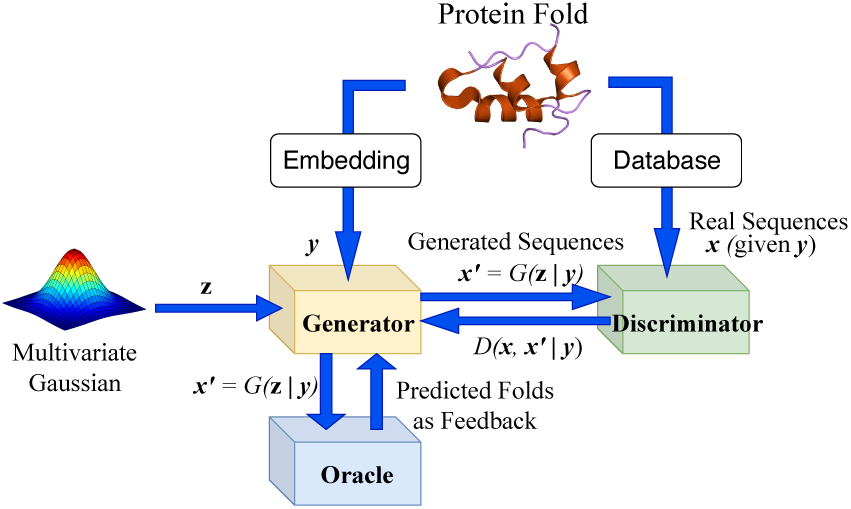
The architecture of guided conditional Wasserstein GAN.

## 2 Materials and Methods

### 2.1 Data

#### 2.1.1 Pre-processing

Sequences labeled with structural folds are retrieved from SCOPe v. 2.07 (Chandonia *et al*., 2018) and filtered at 100% identity level. After processing, the labeled data consists of 20,125 sequences (over 35% of the original) labeled with 781 of the original 1,232 folds, across all 7 fold classes (a—g) and 3 “difficulty” levels based on sequence abundance (at least 50 sequences for easy, at most 5 for hard, and in between for medium). More details are provided in Supporting Information (SI) Sec. S1.

The resulting labeled dataset is split into training (70%), validation (15%) and test (15%) sets with stratified sampling to preserve the fold-class distribution. Folds do not overlap among sets and their statistics are in Table S2. The training sequence statistics are in Table S3. Sequence and structure representatives of each fold are chosen for post-analysis (Sec. S2).

In addition, 31,961 unlabeled sequences without paired structures are randomly obtained from UniRef50 (Suzek *et al*., 2014).

### 2.2 Fold Representation

We aim at a low-dimensional representation of the fold space that are 1) representative enough for preserving the information about the known folds and 2) generalizable enough for describing a novel fold. Considering that the growth of the fold space in recent years has been slow and likely near saturation (Jaroszewski *et al*., 2009), we focus on the space spanned by the 1,232 basis folds from SCOPe v. 2.07 and perform dimension reduction using kernel principal component analysis (Schölkopf *et al*., 1997). This approach is inspired by (Hou *et al*., 2003). Any fold with a structure can be represented with coordinates in the space spanned by the top *K* eigenvectors. More details can be found in SI Sec. S2.

### 2.3 Oracle for Protein Fold Prediction

An oracle predicting protein folds from sequences is used both for guiding sequence generation during model training and for filtering generated sequences once the model is trained. Our oracle is a revised model based on DeepSF (Hou *et al*., 2017). It uses less features for speed and more advanced network architecture for accuracy compared to DeepSF.

First, DeepSF uses input features including amino-acid sequence, position-specific scoring matrix (PSSM), predicted secondary structure (SS), and predicted solvent accessibility (SA); whereas our oracle only uses sequence. The features other than sequences are very informative for fold prediction but unfortunately require computationally expensive multiple sequence alignment. Considering that our model for protein fold design involves millions of generated sequences during training, online calculation of non-sequence features for oracle feedback is infeasible (each sequence demands tens of minutes for non-sequence features). We therefore only use sequence as input feature (one-hot encoding).

Second, DeepSF involves a 1D deep convolutional neural network whereas our oracle is deeper with 10 more layers of residual convolutional layers and its filter size is higher (40 versus 10). The architecture change is to compensate the loss of informative non-sequence features. In the end, a softmax layer predicts the probability for each of 1,215 folds (slightly increased from 1,195 in DeepSF due to SCOPe update). More details including illustration of the oracle architecture can be found in Sec. S3.

### 2.4 GAN Models for De Novo Protein Design

#### 2.4.1 Conditional Wasserstein GAN (cWGAN)

Generative Adversarial Network (GAN) (Goodfellow *et al*., 2014), a class of generative models, represents a game between a generator *G* and a discriminator *D*. The generator’s objective is to generate artificial data from a noise input that are close to real data and the discriminator’s goal is to discriminates the generated data from the real ones.

Compared to the original GAN (Goodfellow *et al*., 2014), Wasserstein GAN (WGAN) (Arjovsky *et al*., 2017; Gulrajani *et al*., 2017) uses Wasserstein distance rather than KL-divergence. This change overcomes training difficulties in GAN, such as difficulty to reach Nash equilibrium (Salimans *et al*., 2016), low dimensional support, and vanishing gradients and mode collapsing (Arjovsky *et al*., 2017).

We consider conditional WGAN (cWGAN) (formulated using the Kantorovich-Rubinstein duality) with gradient penalty, a soft version of Lipschitz constraint modeled through the penalty on the norm of the gradient (Gulrajani *et al*., 2017). The formulation is as follows:

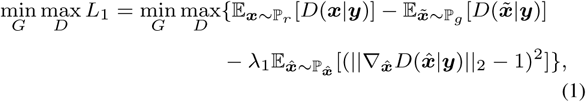

where ***y*** is the embedding of protein fold explained in Sec. 2.2; ***x*** denotes real sequences generated from ℙ_*r*_, the real data distribution; and 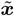 denotes artificial s equences g enerated from ℙ_*g*_, t he m odel d istribution. ℙ_*g*_ is implicitly defined by distribution *p*(**z**) of noise ***z*** because 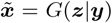. Moreover 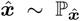 is implicitly defined a s u niformly s ampling along straight lines between pairs of points sampled from ℙ_*g*_ and ℙ_*r*_ for a given label ***y***. This is inspired by the fact that the optimal critic contains straight lines, with l2 norm of the gradient equal to 1, connecting coupled points from ℙ_*g*_ and ℙ_*r*_. Hyper-parameter *λ*_1_ enforces the importance of gradient penalty in the loss function and is set at 10 (Gulrajani *et al*., 2017) without optimization. Pseudo-code of cWGAN with gradient penalty is given in Algorithm 1 in SI.

#### 2.4.2 Semi-supervised cWGAN

When using cWGAN to generate protein sequences for a specific structural fold, a fundamental question remains about whether the sequences are “protein-like” (for instance, whether these sequences could fold into stable and functional structures). To this end, we have further developed a semi-supervised cWGAN by exploiting abundant protein sequences without paired structure data. Specifically, we first train the cWGAN model using unsupervised protein sequences from UniRef50 while fixing their ***y*** at the center of all fold representations. Then we re-train the cWGAN model using the supervised sequences (Sec. 2.1) with corresponding fold representations (Sec. 2.2) but initialize model parameters at the optimal values from the unsupervised step (warm start). The two-step semi-supervised setting could help improve the chance of generating relevant protein sequences and overcome the instability of cWGAN training.

#### 2.4.3 Guided cWGAN (gcWGAN)

In principle, cWGAN could guide the sequence generation specifically for a desired fold. However, limited resolution of fold embedding could present a barrier. We have thus developed a novel GAN model, guided cWGAN (or gcWGAN), with an additional oracle inside. The oracle provides feedback to the generator on sequences generated. Specifically, this feedback is introduced as an additional “regularization” term *R* to *L*_1_ in the loss function:

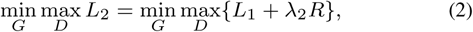

where the hyper-parameter *λ*_2_ is used to balance the two terms.

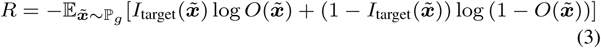

where 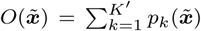, the sum of probabilities for the top *K*′ (10 in this study) fold predictions from the oracle. Ideally, 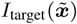 is an indicator function that equals 1 when the target fold is among the top *K*′ predictions from the oracle. But this definition would lead to a non-differentiable expression without gradients needed for back propagation.

We thus introduce 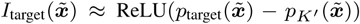 and 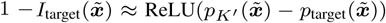. So if the target fold is within top *K*′ predictions, 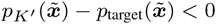 and its RELU assigns zero to 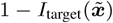; otherwise, zero is assigned to 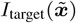.

In guided cWGAN, the oracle is pre-trained as explained in Sec. 2.3 and then fixed, and the generator and the discriminator (more often referred to as the critic in WGAN) are trained with warm start (using optimal parameters from semi-supervised cWGAN in Sec. 2.4.2). Pseudo-code of guided cWGAN is given in Algorithm 2 in SI. Deep learning architectures of the oracle, the critic, and the generator are detailed in Sec. S4.3.

### 2.5 Hyperparameter Tuning

While training cWGAN or gcWGAN, we consider three common hyper-parameters: 1) the initial learning rate for the Adam optimizer, 2) the number of critic iterations, and 3) the noise length. Assuming that optimal hyper-parameters are similar between cWGAN and gcWGAN, we sequentially tune them for cWGAN by training cWGAN for 100 epochs. For gcWGAN, the three common hyper-parameters are adopted the optimal values for cWGAN and its *λ*_2_ is tuned further.

To tune the hyper-parameters, we select four criteria for the intermediate assessment of sequences at each epoch. The criteria are of increasing biological relevance: 1) mathematical convergence through the critic’s loss, 2) low ratio of the “nonsense” sequences (how often padding characters appears between or in front of amino-acid characters to produce invalid sequences), 3) low ratio of padding characters at the end of valid sequences, and 4) sequence novelty through low sequence identity between generated sequences and real representative sequences for a given fold.

More details are provided in Sec. S5.

### 2.6 Assessment

Once the generator is trained as part of gcWGAN, we feed it with the desired structural fold (in its embedding ***y*** as described in Sec. 2.2) and generate sequences to pass the nonsense check and then the oracle’s check. The intermediate, valid sequences (before the final, oracle’s check) are generated up to 10^5^ per fold and assessed across all folds using the oracle, a fold classifier. The final sequences (after passing the oracle) are generated 10 per fold and assessed over selected representative folds using Rosetta, an *ab initio* structure predictor.

#### 2.6.1 Assessing Intermediate Designs

Once the generator is trained, we first examine “yield ratios” of generated valid sequences (passing the nonsense check). Specifically, a generated sequence is declared a “yield” if the target fold is within the top-10 folds predicted by the oracle; and the yield ratio is the portion of those yields. When the target fold is a novel one not defined for the oracle, we use its neighbors within 0.3 distance unit in the embedded fold space (0.3 is chosen using the training set so that each fold has about 3 neighbors on average). Starting with 10^3^ valid sequences per fold, we incrementally generate up to 10^5^ sequences at a stepsize of 10^3^, until finding 10 yields.

#### 2.6.2 Assessing Final Designs

We further analyze the fold accuracy of the final designs (those yields) by predicting their structures using Rosetta (Leaver-Fay *et al*., 2011). Specifically, for each target fold, we generate 100 valid sequences that pass the final check of the oracle and “fold” the first ten such sequences into 100 structure predictions using Rosetta. We then evaluate their structural accuracy compared to the known representative structure using TM score. We also evaluate sequence diversity among each set of 100 sequences and their sequence novelty compared to the known representative sequence.

##### Structure Prediction

We perform *ab initio* structure prediction using Rosetta v. 3.10 for each sequence. Specifically, we generate 10,000 trajectories the 10,000 structural predictions, we retain around 10% energetically-lowest ones by applying an energy-cutoff of 200, cluster those with a cluster radius of 2.5Å in RMS and a maximum cluster count of 10, and report the 10 cluster centers as final structure predictions.

##### Structure Accuracy

We align each structure prediction of a designed sequence to the known, representative structure of its target fold, using TM-align (Zhang and Skolnick, 2005); and calculate its TM-score (the reference being the target structure). A TM-score, between 0 and 1, indicates essentially the same fold when it is above 0.5 and non-random similarity when it is above 0.3. We also calculate the normalized ΔTM-score by subtracting the TM-score with a background value. The background value is simply the average TM-score between the predicted structure and representative structures of all off-target folds. A positive ΔTM-score would indicate a sequence design specific to the target fold.

##### Sequence Diversity and Novelty

We perform pairwise alignment of the 100 final sequences for each design and calculate the distribution of sequence identity to measure diversity. We also do so between each designed sequence and the known, representative sequence for each target fold, to measure sequence novelty. As a sequence identity value above 0.3 would indicate close homologs whose structures are very likely similar, we regard 0.3 as a threshold below which generated sequences are diverse or novel.

##### Selected Folds

Since structure prediction is computationally expensive (10^4^ core-hours per sequence), we choose six representative test folds across fold classes (a–d and g), sequence abundance (easy to hard, reflecting “designability”), and yield ratios (above 0.01 for high and otherwise for low):

In addition, to check the model performance for prospective, novel folds, we also select a recently published fold ElGamacy *et al*. (2018) that does not belong to the 1,232 basis folds.

## 3 Results

### 3.1 Fold Representation

To construct a low-dimensional fold representation, we performed kernel PCA for the fold space spanned by the 1,232 basis folds. The first 20, 200, and 400 principal components explained around 17%, 50%, and 75% of the variance (Fig. S2). As the dimension of fold representation determines that of the conditioning variable ***y*** in cWGAN and a higher dimension causes more demanding model training, we chose the space spanned by the first 20 principal components as a lower-dimensional representation of fold space. An analysis on the resolution of the fold representation shows that the explained variance of the 20 principal components reach 63% for 100 cluster centers of the 1,232 original folds (Fig. S3). Visualization of the fold representations shows that folds are well-clustered consistent with their class membership (Fig. S4), where *α*/*β* and *α*+*β* folds’ representations distributed between the clusters of *α* and *β* folds.

### 3.2 Oracle’s Accuracy

To assess the oracle as a fold predictor, we compared, in Table S4, the original DeepSF using all features or sequence only and our oracle (modified DeepSF) using sequence only. For the test set, top-10 predictions from DeepSF impressively achieved an accuracy of 0.94 using all features; whereas they only did that of 0.69 using just sequence. By modifying the model architecture, our oracle using just sequence increased the accuracy to 0.74. Meanwhile, our oracle only uses milliseconds to predict for each sequence whereas DeepSF does minutes on non-sequence feature calculation alone. Therefore, our oracle is a somewhat ambiguous yet ultra-fast fold predictor that is suitable for the framework of gcWGAN.

### 3.3 Semi-Supervision Improves Training for gcWGAN

We report hyper-parameter tuning results in Fig. S6–S8 to showcase the benefit of semi-supervision. The training of gcWGAN was warmed up with parameters initialized at trained values of the semi-supervised cWGAN. How does it compare against the scenario without semi-supervision, i.e., when the parameters were initialized at cWGAN values rather than the semi-supervised cWGAN ones. Semi-supervised gcWGAN had lower overall loss in the last 30 epochs, mostly due to lower critic losses (Fig. S9). Therefore, semi-supervision improved the convergence of gcWGAN model training.

### 3.4 Oracle Feedback Increases Yields

For intermediate sequence designs, we first examine their yield ratios for all structural folds in Table 2. gcWGAN with oracle feedback improved the yield ratio for an average test fold by around 15% compared to cWGAN; did so for 4 out of 7 fold classes; and did so by over 80% for difficult cases that are the least designable. Two factors affect the yield ratios: (1) easy folds with abundant sequence availability are with higher yield ratios for training or test set because of more data or more designability; (2) folds for which the oracle is more accurate see higher yield ratios (Table S5).

**Table 1.**
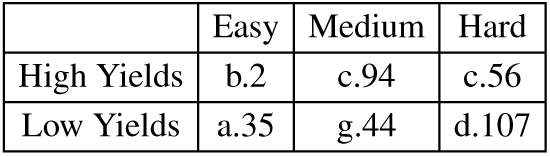
Representative test folds for structure assessment.

|  | Easy | Medium | Hard |
| --- | --- | --- | --- |
| High Yields | b.2 | c.94 | c.56 |
| Low Yields | a.35 | g.44 | d.107 |

**Table 2.**
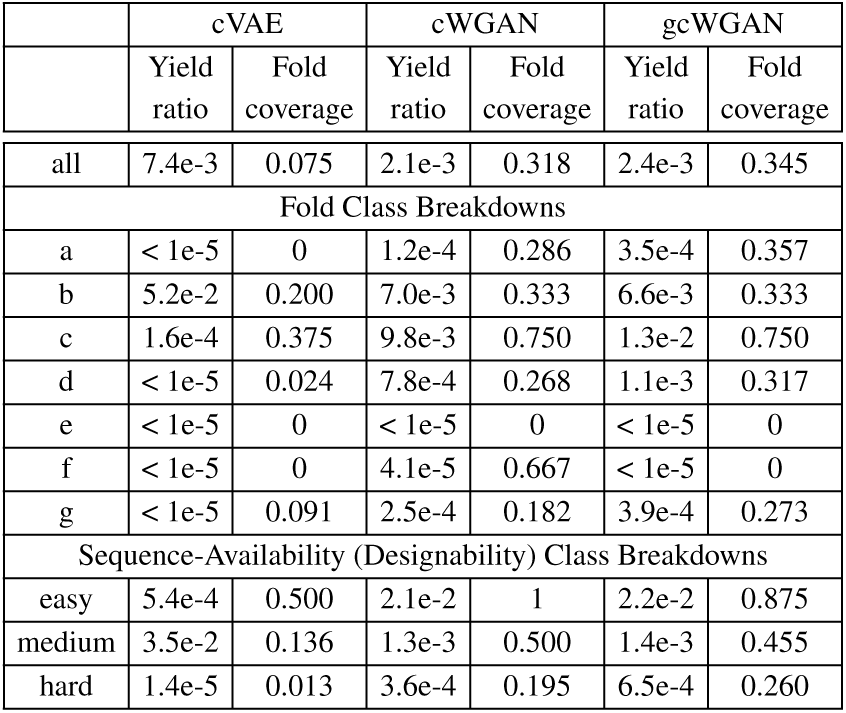
Yields among cVAE, cWGAN and gcWGAN over test folds.

We also incrementally generate 10^5^ sequences for each of the six selected folds. We observe that gcWGAN with feedback increased the yield efficiency by 18% to 205% for five of them (Fig. S10). For the sixth fold (d.107), gcWGAN produced a yield whereas cWGAN failed to.

### 3.5 gcWGAN Improves Yield Ratios for Most Folds Compared to cVAE

We compared yield ratios between gcWGAN and a recent cVAE-based software for protein design (Greener *et al*., 2018) in Table 2 (all folds and breakdowns) and S7 (six selected folds). Overall, gcWGAN had higher yield ratios for four of seven fold classes and comparable ones for another two. The average yield ratio is higher with cVAE, which is very misleading because the average for cVAE is dominated by an extremely high yield ratio for only one fold. Importantly, gcWGAN achieved yield ratios over 1E-5 for 34.5% of test folds whereas cVAE only did 7.5%. For all six selected test folds, gcWGAN increased the yield ratios by 1 to 2 orders of magnitude. Note that these test folds cannot be ruled out as possible training folds for the published cVAE model due to the lack of information.

All the yield ratios are relatively low – gcWGAN achieved around 2% on average even for folds with abundant sequences or high oracle-accuracy. We however note that many sequences declared non-yields by the imprecise oracle could be false negatives. We also note that low yield ratios can be overcome with high throughput and do not affect accuracy as we will show next.

### 3.6 gcWGAN Designs Sequences of Comparable or Better Accuracy Compared to cVAE

We proceed to examine final sequence designs (the yields) using Rosetta-based structure prediction, for six selected test folds (not seen in the training set thus regarded novel). For each fold, we designed 10 sequences using either gcWGAN and cVAE and predicted 100 structure models using Rosetta. The distributions of the structures’ TM-scores (Fig. 2), when compared to corresponding ground truth, show that gcWGAN outperform cVAE. Specifically, gcWGAN had higher TM-score distributions than cVAE for all six folds except g.44 (comparable) with p-values way below 5E-2 (Table S8). Similarly, gcWGAN had better ΔTM-score distributions for three folds and comparable ones for the other three (Fig. S12). Taken together, gcWGAN could design protein sequences more specific to target folds than off-targets. And it can sometimes do so with good accuracy: the best TM-score is above 0.5, indicating the same fold, for fold a.35; and the score is below 0.5 but above 0.4 for two other folds. The best sequence/structure combination for each fold is superimposed to the ground truth in Fig. S14.

**Fig. 2:**
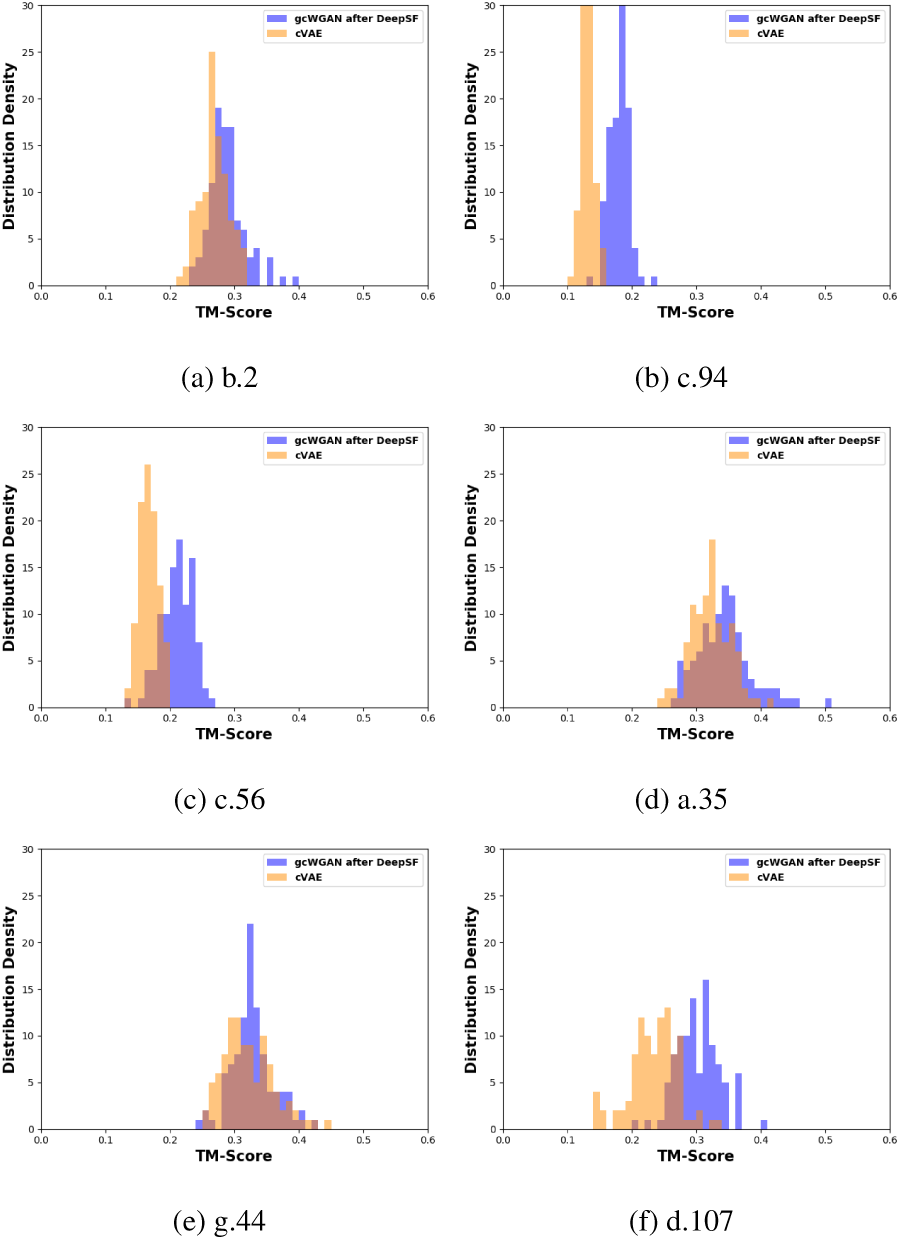
Distribution of the TM scores between cVAE/gcWGAN designs and the ground truth on 6 selected, representative test folds.

We also compared gcWGAN and cVAE on a novel fold not even in the 1,232 basis folds for fold embedding. Again, gcWGAN’s designs achieved better TM-scores and ΔTM-scores than cVAE with statistical significance (p-value below 2E-13 and 0.035, respectively; see Table S8). Impressively, the best sequence/structure combination from gcWGAN had a TM-score of 0.47, very close to the threshold to indicate the same fold. The structure is superimposed to the ground truth in Fig. 3b where 3–4 of 5 helices in ground truth are (partially) aligned to designed helical regions.

**Fig. 3:**
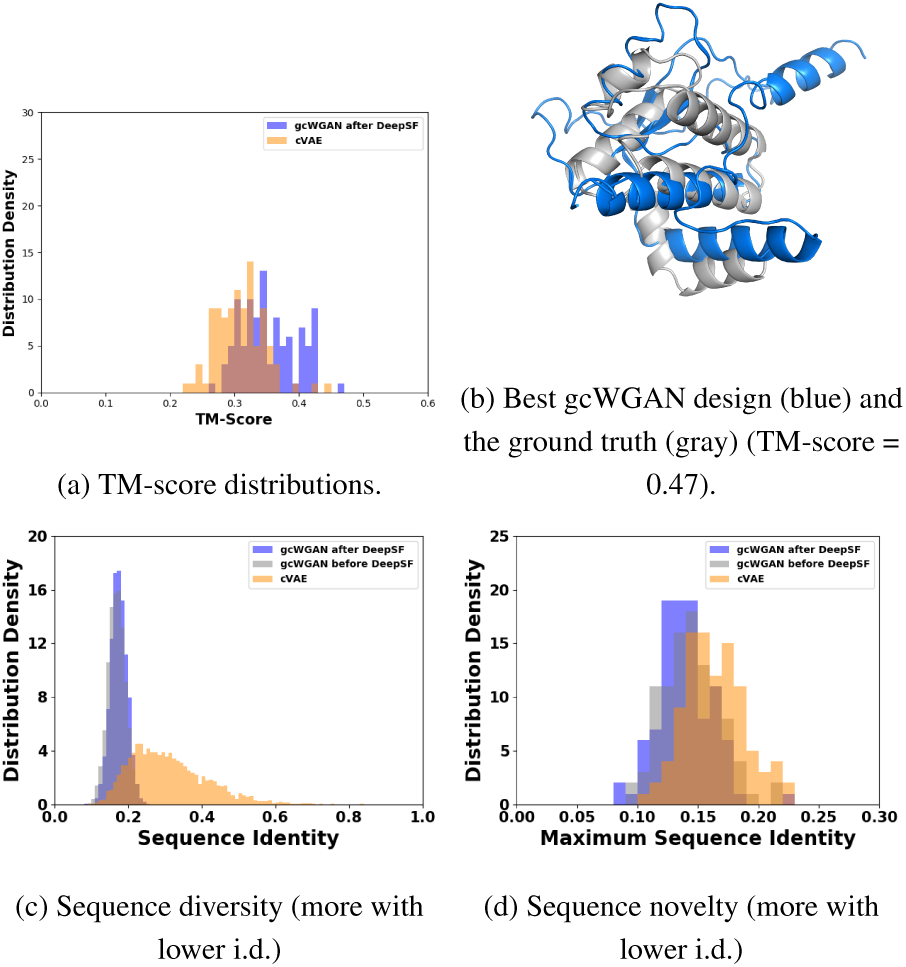
Assesing gcWGAN designs targeting a completely novel fold.

### 3.7 gcWGAN Designs More Diverse and More Novel Sequences Compared to cVAE

Besides the accuracy, we also examine the diversity among the designed protein sequences as well as their novelty compared to known representative sequences. For both the six test folds and the completely novel fold, gcWGAN designs for the same fold are of low sequence identity below 0.3 whereas most cVAE designs are close homologs with sequence identity above 0.3 (Fig. 4 and Fig. 3c). Apparently, gcWGAN could explore much more diverse regions of the sequence space, while maintaining decent fold specificity and accuracy as examined earlier.

**Fig. 4:**
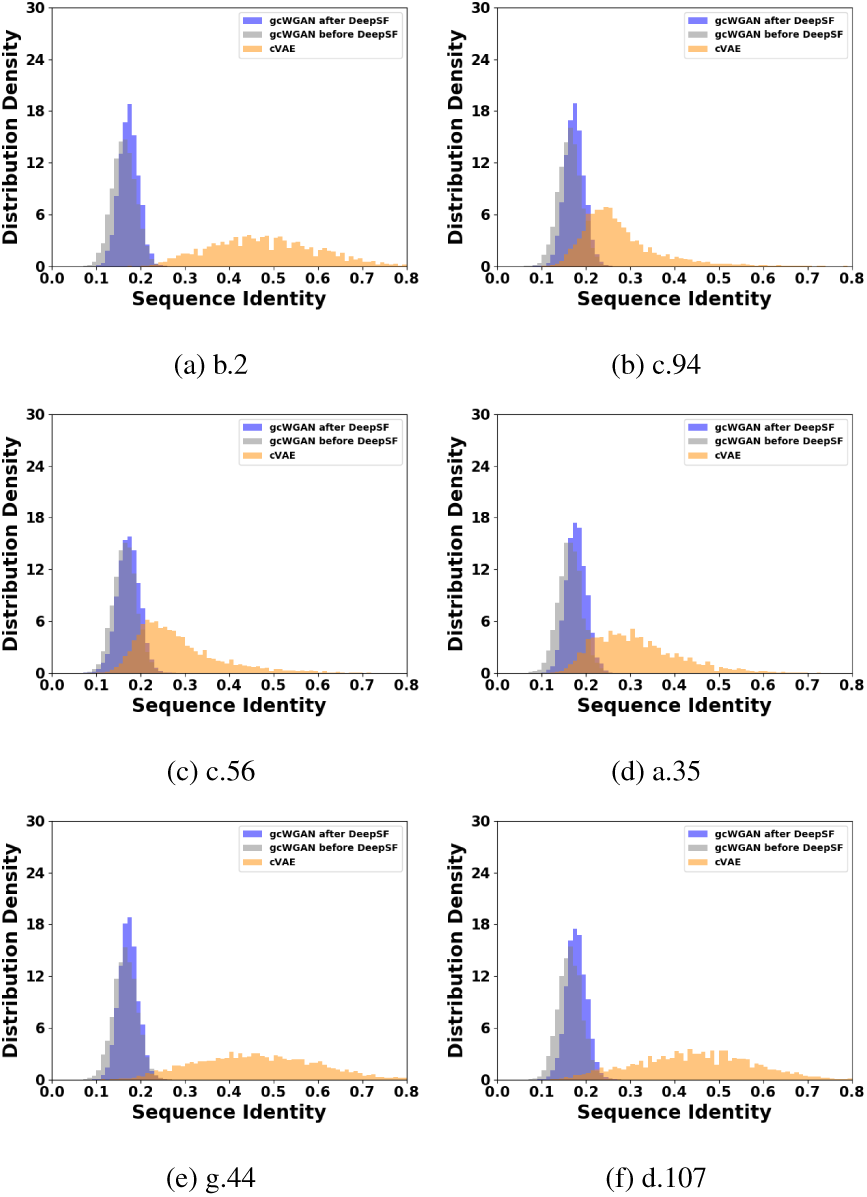
Comparing sequence diversity between cVAE and gcWGAN designs for the 6 selected cases. Lower sequence identity indicates more diversity and that below 0.3 indicates no close homologs.

Moreover, gcWGAN designs are almost entirely of sequence identity lower than 0.3 compared to the closest, known representative sequence for the desired fold (Fig. 5 and Fig. 3d). In contrast, cVAE designs are often close homologs of a known representative sequence with sequence identity above 0.3. Therefore, gcWGAN is exploring uncharted regions in the sequence space.

**Fig. 5:**
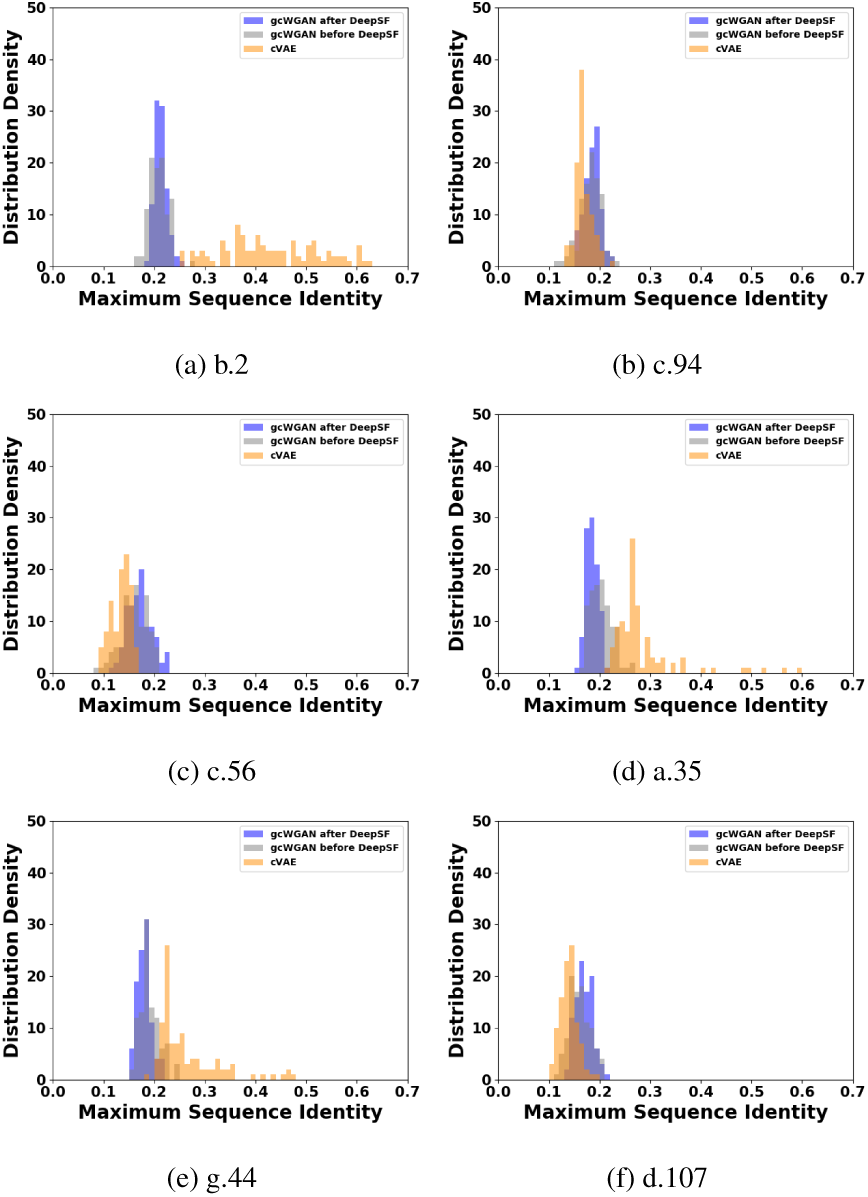
Comparing sequence novelty between cVAE and gcWGAN designs for the 6 selected cases. Lower sequence identity indicates more novelty compared to known representative sequences.

## 4 Conclusions and Discussion

We have designed novel deep generative models for *de novo* protein design targeting novel structure folds. Here we utilize both sequence data with and without structures in a semi-supervised setting and an ultra-fast fast oracle for fold recognition as feedback. As summarized below, our results reveal the value of current protein data toward unraveling sequence-structure relationship and utilizing resulting knowledge for inverse protein design. Over model-designed sequences for nearly 800 folds of diverse class and difficulty (including over 100 test folds not seen in the training set), we first globally assess their performances using the oracle, an ultra-fast yet imprecise fold predictor. Compared to a data-driven alternative (cVAE), our gcWGAN model generates more yields (according to the oracle) for nearly all target folds and achieves much more coverage of target folds. gcWGAN designs proteins much faster than physics-driven structure-based methods (10^−3^s versus at least 10^5^ core hours); thus can tailor a much reduced design space for the latter.

For selected representative test folds and a novel fold not seen in the 1,232 basis folds, we assess gcWGAN designs using a structure predictor. We find that gcWGAN designs are comparable or better in fold accuracy compared to cVAE. Notably, gcWGAN designs are much more diverse, which can be attributed to that WGAN with its implicit model and Wasserstein distance overcomes known limitations of VAE (Yin and Zhou, 2018; Arjovsky *et al*., 2017; Davidson *et al*., 2018). Moreover, gcWGAN designs are often novel in sequence rather than close homologs of known proteins. This indicates that gcWGAN is exploring uncharted regions in the sequence space, using what it learns from the data about sequence-fold relationship rather than simply mimicking the data.

Among major factors affecting gcWGAN performances, fold representation and oracle’s accuracy need the most improvement. Currently, our fold representation is learned in a non-parametric way (kPCA) preserving distance metrics; but the resolution is limited as we choose a low-dimensional subspace for computational concerns. Our oracle is ultra-fast yet highly ambiguous and imprecise, so the feedback has limited performance boost. Interestingly, folds with few sequences known in nature, thus potentially less designable, also appear to be more difficult for the data-driven gcWGAN approach even though those sequences and folds are never known to gcWGAN during training.

## Acknowledgements

Portions of this research were conducted with high performance research computing resources provided by Texas A&M University.

