## Supplementary Information for "De Novo Protein Design for Novel Folds using Guided Conditional Wasserstein Generative Adversarial Networks (gcWGAN)"

### 1 Data

#### 1.1 Pre-processing

We download fold data (including corresponding sequences and structure domains) from SCOPe (v. 2.07) and filtered sequences at 100% identity level. Some uncommon sequences (<2%) contain non-standard amino acid letters including 'b', 'z', 'x' and 'X'. The first three represent the ambiguity among ['d', 'n'], ['q','e'] and all 20 amino acids, respectively; and 'X' represents a gap in sequence. For simplicity, we assign explicit standard amino acids to each occasion of 'b', 'z', and 'x' following a uniform prior and disregard sequences containing 'X'.

To save the computational cost under limited budget, we further filter the remaining and retained those of sequence length between 60 and 160. The length interval is chosen to balance the range (for sequence padding concerns) and the fold space coverage. Specifically, a tight range around the mode of the distribution would lead to sequences of similar lengths and reduce padding needs. Meanwhile, the range needs to be wide enough to cover the sequence and the fold space. In the end, the chosen range represents over 35% of the filtered sequences (Fig. S1) and over 63% of the folds (781 out of 1,232).

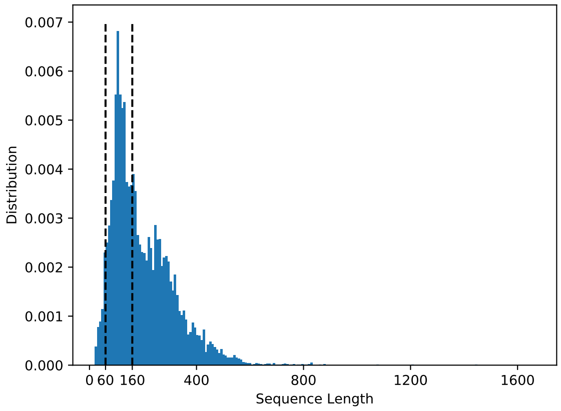


Figure S1: Sequence length distribution in the SCOPe data (100% sequence identity level). The percentage within the interval [60, 160] is 35.87%.

#### 1.2 Statistics

SCOPe clustered the sequences into 7 classes indexed from a to g with names and counts given in Table S1. We also classify the folds into three difficulty categories based on sequence abundance: easy for those with at least 50 sequences, medium for 6 to 49 sequences, and difficult for at most 5 sequences. The split counts based on the combination of fold class and difficulty are provided in Table S1.

| Class Index | Class Name | Easy | Medium | Difficulty | Total |
|---|---|---|---|---|---|
| a | $\alpha$ | 27 | 64 | 105 | 196 |
| b | $\beta$ | 28 | 57 | 54 | 139 |
| c | $\alpha/\beta$ | 15 | 35 | 18 | 68 |
| d | $\alpha+\beta$ | 38 | 110 | 140 | 288 |
| e | Multi-domain proteins | 0 | 0 | 6 | 6 |
| f | Membrane and cell surface | 2 | 9 | 10 | 21 |
| g | Small proteins | 7 | 17 | 39 | 63 |
| | Total | 117 | 292 | 372 | 781 |

Table S1: The statistic of folds in the whole dataset based on class and difficulty.

We use stratified sampling to split the resulting dataset into training (70%), validation (15%) and test (15%) sets while preserving the fold-class distributions.

| Class Index | Class Name | Training Set | Val. Set | Test Set | Total |
|---|---|---|---|---|---|
| a | $\alpha$ | 139 | 29 | 28 | 196 |
| b | $\beta$ | 109 | 15 | 15 | 139 |
| c | $\alpha/\beta$ | 52 | 8 | 8 | 68 |
| d | $\alpha+\beta$ | 206 | 41 | 41 | 288 |
| e | Multi-domain proteins | 3 | 2 | 1 | 6 |
| f | Membrane and cell surface | 14 | 4 | 3 | 21 |
| g | Small proteins | 40 | 12 | 11 | 63 |
| | Total | 563 | 111 | 107 | 781 |

Table S2: The statistics of folds in training, validation and testing sets based on class.

| Class Index | Class Name | # of Seq. |
|---|---|---|
| a | $\alpha$ | 5486 |
| b | $\beta$ | 4562 |
| c | $\alpha/\beta$ | 2536 |
| d | $\alpha+\beta$ | 6625 |
| e | Multi-domain proteins | 8 |
| f | Membrane and cell surface | 189 |
| g | Small proteins | 719 |
| | Total | 20125 |

Table S3: The statistics of sequences in training set.

Sequence representative(s) of each fold are chosen for post-analysis. With pairwise sequence identities as similarity measure, all sequences of the same fold are clustered using DBSCAN (Ester *et al.*, 1996) ($\epsilon = 0.7$ as proteins of sequence identity over 0.3 very likely have similar structures). A representative is chosen for each cluster based on the maximum sum of sequence identities towards all other sequences within the same cluster.
A single structure representative is already chosen for each fold by SCOPe.

### 2 Fold representation

#### 2.1 Methods

Using a representative protein structure chosen by SCOPe for each of the 1,232 folds, we construct a pairwise similarity matrix of symmetrized TM scores (Zhang and Skolnick, 2004) and added a properly-scaled identity matrix to it to make a positive-definite Gram matrix $\boldsymbol{S}$. We centralize the Gram matrix $\boldsymbol{S}$ through:

$$\begin{aligned} \boldsymbol{O}_{\mathrm{NN}} &= \mathbf{1}_{\mathrm{N}}\mathbf{1}_{\mathrm{N}}^{T}/N; \\ \tilde{\boldsymbol{S}} &= (\boldsymbol{I} - \boldsymbol{O}_{\mathrm{NN}})\boldsymbol{S}(\boldsymbol{I} - \boldsymbol{O}_{\mathrm{NN}}); \end{aligned} \tag{1}$$

where $N$ being 1,232 is the row and column size of $\boldsymbol{S}$, $\mathbf{1}_{\mathrm{N}}$ is an all-one column vector of size $N$ by 1, and $\boldsymbol{I}$ is an identity matrix of size $N$ by $N$. We then perform eigen-decomposition to $\tilde{\boldsymbol{S}}$ and rank its eigenvalues $u_i$ and its corresponding eigenvectors $\boldsymbol{v}_i$ in a decreasing order: $\{(u_1, \boldsymbol{v}_1), (u_2, \boldsymbol{v}_2), ..., (u_N, \boldsymbol{v}_N)\}$. The $i$th dimension basis of fold space is calculated through:

$$\boldsymbol{r}_i = \frac{\boldsymbol{v}_i}{\sqrt{u_i}} \tag{2}$$

Then for $M$ incoming new folds, we first calculate their TM scores with individual $N$ basis folds and form a new matrix $\boldsymbol{S}^*$ with size $M \times N$. We then centralize $\boldsymbol{S}^*$ though

$$\begin{aligned} \boldsymbol{O}_{\mathrm{MN}} &= \mathbf{1}_{\mathrm{M}}\mathbf{1}_{\mathrm{N}}^{T}/N; \\ \tilde{\boldsymbol{S}}^* &= \boldsymbol{S}^* - \boldsymbol{S}^*\boldsymbol{O}_{\mathrm{NN}} - \boldsymbol{O}_{\mathrm{MN}}\boldsymbol{S} + \boldsymbol{O}_{\mathrm{MN}}\boldsymbol{S}\boldsymbol{O}_{\mathrm{NN}}; \end{aligned} \tag{3}$$

And then the coordinate vector of these these $M$ folds along the $i$th dimension is $\tilde{\boldsymbol{S}}^* \cdot \boldsymbol{r}_i$.

#### 2.2 Results

We first examine how much of the variance in the original fold space spanned by 1,232 basis folds can be explained by the top eigenvectors from kernel PCA (kPCA). As seen in Fig. S2, the first 20 eigenvectors explained 17.3% of the variance in the original fold space spanned by 1,232 basis folds.

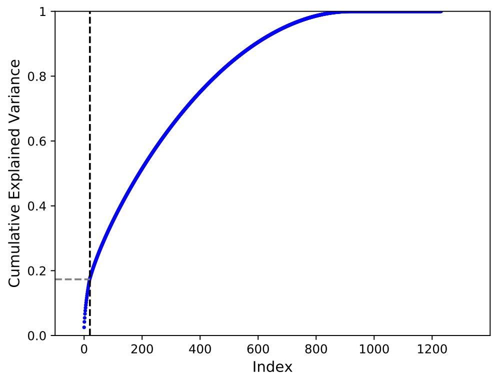


Figure S2: Cumulative variance explained by the top eigenvectors of kernel PCA for fold representation.

Due to the computational limit of deep generative models, we have to restrict the fold representation to a space spanned by the 20 eigenvectors. In this case, we ask what resolution of the fold space is accomplished by the representation. To answer the question, we cluster these 1,232 folds into varying $K$ clusters through K-means clustering. The centriods of these $K$ clusters form a $K$-dimensional subspace for the original 1,232-dimensional space. We then project the aforementioned 20 eigenvectors into this subspace and calculate the explained variance of those 20 projected vectors for the subspace. As seen in Fig. S3, the 20 eigenvectors could explain over 63% of the variance in a fold subspace spanned by 100 representative folds.

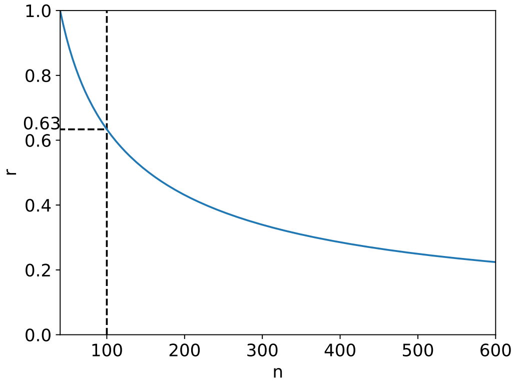


Figure S3: The explained variance of the $K$ clusters (or the $K$D subspace of the 1232D original fold space) using the aforementioned 20 eigenvectors.

The new representations of 1,232 folds are visualized in the a 3D space spanned by the first 3 of the 20 principal components in Fig. S4.

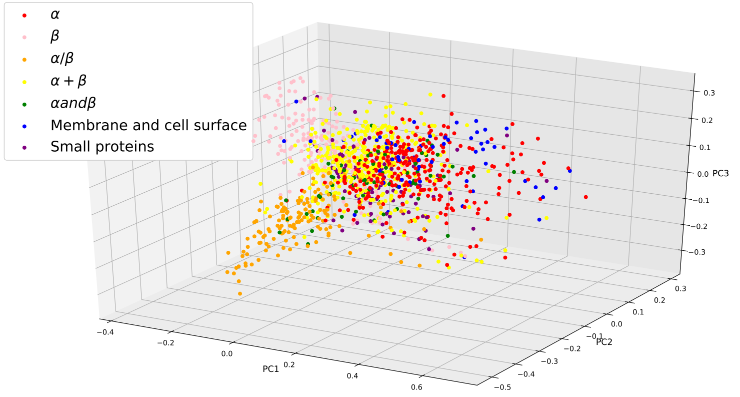


Figure S4: Visualization of 1,232 folds in a space spanned by the top 3 principal components found by kPCA.

### 3 Oracle for Protein Fold Prediction

#### 3.1 Methods

An oracle of protein fold recognition for a query sequence is needed for both guiding sequence generation during model training and assessing accuracy once the model is trained. Since there is no closed-form mathematical formula for protein fold as a function of the sequence, protein fold is often predicted through data-driven approaches such as DeepSF (Hou *et al.*, 2017). Hou and co-workers designed a 1D deep convolutional neural network to classify protein folds with variable-length to each of 1,195 protein folds. The input features include 1) amino acid (AA) sequences, 2) position-specific scoring matrix (PSSM), 3) predicted secondary structure (SS), and 4) predicted solvent accessibility (SA). PSI-BLAST for multiple sequence alignment is required for the calculation of PSSM, SS and SA. PSI-BLAST costs minutes for each sequence and can be tolerated for one-time feature generation for few sequences. However, it is computationally daunting for on-line feature calculation for the huge number of sequences generated during our model training.

Therefore, we modified DeepSF to classify a protein into 1,215 folds (slightly increased to consider SCOPe update) using one-hot encoding of sequence alone. Since sequence features were found the least informative, such a model would significantly suffer in accuracy. We thus further modified the architecture of DeepSF neural networks to be wider and deeper (with residual blocks). Specifically, we increase the filter size from 10 to 40; and deepen the model from 10 layers of 1D convolution to 20 layers by inserting 10 residual convolutional layers. Modified DeepSF architecture is shown as part of Fig. S5.
Modified DeepSF consists of two branches with filter sizes of 6 and 10 to capture short and long motif information from the sequence, respectively. Each branch consists of blocks of convolution layers followed by batch normalization and ReLU activation. These blocks are non-residual at the beginning and at the end, but residual (connected through skip connections) in the middle. At the end of each branch, a customized max pooling layer has been implemented to capture only the top $m$ amino acids from the sequence. This max pooling is required to deal with the length varying sequences so that its output is of fixed length $m$ (In our model $m$ is equal to 30). Lastly, considering the limited accuracy of the top-1 fold predictions, we further allow the ambiguity of the oracle and use its top-10 predictions to guide or assess each generated sequence.

#### 3.2 Results

We compare the accuracy for fold prediction between the original and the modified DeepSF in Table S4. The model trained for the 1,215 folds using sequence only is adopted as the oracle.

| Prediction | Training set | | | | Test set | | | |
|---|---|---|---|---|---|---|---|---|
| | Original DeepSF | | Modified DeepSF | | Original DeepSF | | Modified DeepSF | |
| | All features (1,195 folds) | AA only (1,195 folds) | AA only (1,195 folds) | AA only (1,215 folds) | All features (1,195 folds) | AA only (1,195 folds) | AA only (1,195 folds) | AA only (1,215 folds) |
| Top1 | 0.74 | 0.36 | 0.56 | 0.49 | 0.72 | 0.33 | 0.40 | 0.36 |
| Top5 | 0.93 | 0.65 | 0.82 | 0.75 | 0.9 | 0.59 | 0.63 | 0.61 |
| Top10 | 0.97 | 0.76 | 0.89 | 0.84 | 0.94 | 0.69 | 0.74 | 0.72 |
| Top15 | 0.98 | 0.82 | 0.92 | 0.88 | 0.96 | 0.74 | 0.79 | 0.78 |
| Top20 | 0.98 | 0.86 | 0.94 | 0.91 | 0.97 | 0.79 | 0.82 | 0.81 |

Table S4: Fold prediction accuracy for the original and our modified DeepSF models with various features and architectures.

### 4 GAN Models for De Novo Protein Design

Generative Adversarial Network (GAN) (Goodfellow *et al.*, 2014), a very successful and powerful class of generative models, represents a game between a generator $G$ and a discriminator $D$. The generator's objective is to generates artificial data from a noise input that are close to real data and the discriminator's goal is to discriminates the generated data from the real ones. GAN has wide applications such as generating images of unprecedented quality of celebrities (Karras *et al.*, 2017), image to image translation (Chu *et al.*, 2017), text to image translation (Zhang *et al.*, 2017), creating anime characters (Jin *et al.*, 2017), image inpainting (Pathak *et al.*, 2016), face aging (Antipov *et al.*, 2017) and music generation (Yang *et al.*, 2017). It has also been extended from an unsupervised generative model to a supervised one through conditional GAN (Mirza and Osindero, 2014) that generates images specific for each class.
Despite their popularity, GANs are notoriously hard to train due to problems such as the difficulty to achieve Nash equilibrium (Salimans *et al.*, 2016), low dimensional support, vanishing gradient and mode collapsing (Arjovsky *et al.*, 2017). Wasserstein GAN (WGAN) (Arjovsky *et al.*, 2017; Gulrajani *et al.*, 2017) is a fundamental improvement to the original GANs that addresses these problems. WGAN replaces the KL-divergence with Wasserstein (or earth-mover's) distance. The latter has less issues toward low dimensional support (thus more stable) and use the notation critics instead of the discriminator.

#### 4.1 Conditional WGAN

See **Algorithm 1**

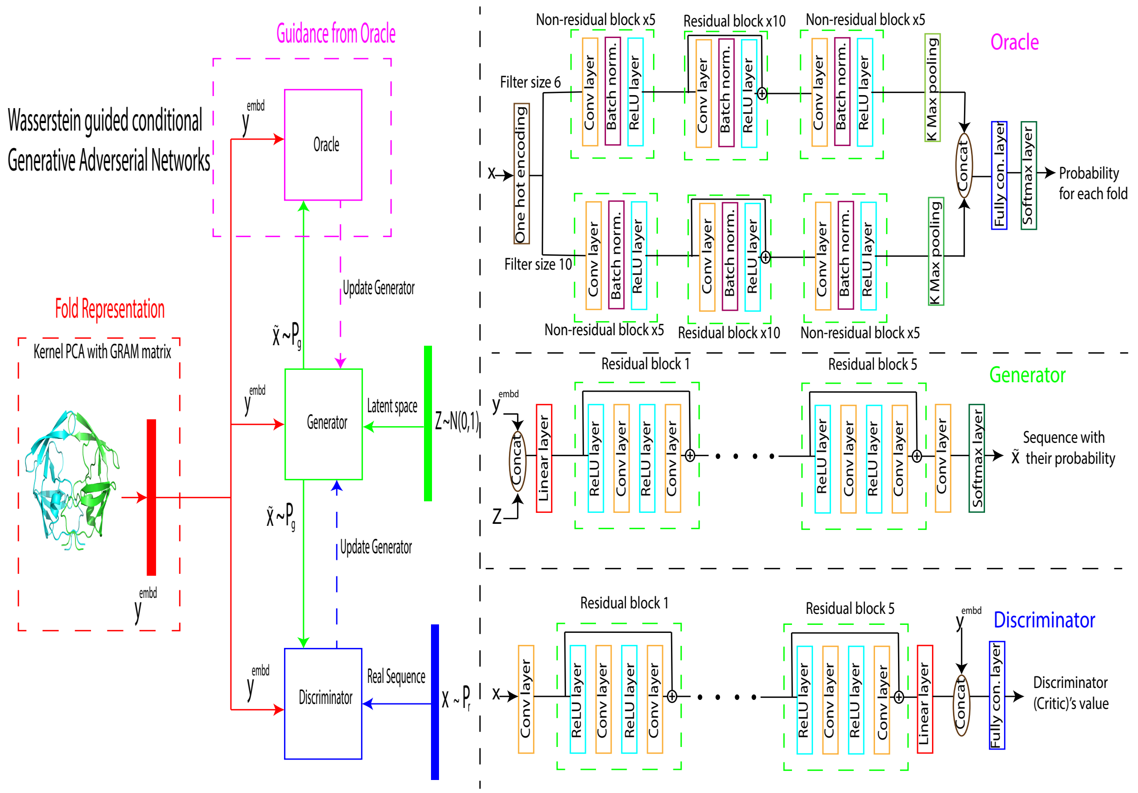


Figure S5: The overall architecture of our guided conditional Wasserstein GAN (gcWGAN) including those of the oracle, the generator, and the discriminator.

**Algorithm 1** Conditional WGAN with gradient penalty

Randomly initialize the parameter from scratch
**while** $\theta$ has not converged **do**
  **for** t=1$\cdots n_{\text{critic}}$ **do**
    Sample real data $\{\mathbf{x}^{(i)}, \mathbf{y}^{(i)}\}_{i=1}^{m} \sim \mathbb{P}_r$
    Sample noise $\{\mathbf{z}^{(i)}\}_{i=1}^{m} \sim p(\mathbf{z})$
    Sample random number $\{\epsilon^{(i)}\}_{i=1}^{m} \sim U[0,1]$
    $\{\tilde{\mathbf{x}}^{(i)}\}_{i=1}^{m} \leftarrow \{G_\theta(\mathbf{z}^{(i)}|\mathbf{y}^{(i)})\}_{i=1}^{m}$
    $\{\hat{\mathbf{x}}^{(i)}\}_{i=1}^{m} \leftarrow \{\epsilon^{(i)}\mathbf{x}^{(i)} + (1-\epsilon^{(i)})\tilde{\mathbf{x}}^{(i)}\}_{i=1}^{m}$
    $L \leftarrow \frac{1}{m}\sum_{i=1}^{m} D_w(\mathbf{x}^{(i)}|\mathbf{y}^{(i)}) - D_w(\tilde{\mathbf{x}}^{(i)}|\mathbf{y}^{(i)}) - \lambda(||\nabla_{\hat{\mathbf{x}}^{(i)}} D_w(\hat{\mathbf{x}}^{(i)}|\mathbf{y}^{(i)})||_2 - 1)^2$
    $w \leftarrow \text{Adam}(\nabla_w L, w, \alpha, \beta_1, \beta_2)$
  Sample a batch of noises $\{\mathbf{z}^{(i)}\}_{i=1}^{m} \sim p(\mathbf{z})$
  $\theta \leftarrow \text{Adam}(\nabla_\theta \frac{1}{m}\sum_{i=1}^{m} -D_w(G_\theta(\mathbf{z}^{(i)}|\mathbf{y}^{(i)})), \theta, \alpha, \beta_1, \beta_2)$

#### 4.2 Guided cWGAN (gcWGAN)

See **Algorithm 2**

**Algorithm 2** guided cWGAN with gradient penalty

Initialize the parameter from the previously trained semi-supervised cWGAN
**while** $\theta$ has not converged **do**
  **for** t=1$\cdots$,$n_{critic}$ **do**
    Sample real data $\{\mathbf{x}^{(i)}, \mathbf{y}^{(i)}\}_{i=1}^{m} \sim \mathbb{P}_r$
    Sample noise $\{\mathbf{z}^{(i)}\}_{i=1}^{m} \sim p(\mathbf{z})$
    Sample random number $\{\epsilon^{(i)}\}_{i=1}^{m} \sim U[0,1]$
    $\{\tilde{\mathbf{x}}^{(i)}\}_{i=1}^{m} \leftarrow \{G_\theta(\mathbf{z}^{(i)}|\mathbf{y}^{(i)})\}_{i=1}^{m}$
    $\{\hat{\mathbf{x}}^{(i)}\}_{i=1}^{m} \leftarrow \{\epsilon^{(i)}\mathbf{x}^{(i)} + (1-\epsilon^{(i)})\tilde{\mathbf{x}}^{(i)}\}_{i=1}^{m}$
    $L_D \leftarrow \frac{1}{m}\sum_{i=1}^{m} D_w(\mathbf{x}^{(i)}|\mathbf{y}^{(i)}) - D_w(\tilde{\mathbf{x}}^{(i)}|\mathbf{y}^{(i)}) - \lambda_1(||\nabla_{\hat{\mathbf{x}}^{(i)}} D_w(\hat{\mathbf{x}}^{(i)}|\mathbf{y}^{(i)})||_2 - 1)^2$
    $w \leftarrow \text{Adam}(\nabla_w L_D, w, \alpha, \beta_1, \beta_2)$
  Sample a batch of noises $\{\mathbf{z}^{(i)}\}_{i=1}^{m} \sim p(\mathbf{z})$
  Sample real data $\{\mathbf{x}^{(i)}, \mathbf{y}^{(i)}\}_{i=1}^{m} \sim \mathbb{P}_r$
  $\{\tilde{\mathbf{x}}^{(i)}\}_{i=1}^{m} \leftarrow \{G_\theta(\mathbf{z}^{(i)}|\mathbf{y}^{(i)})\}_{i=1}^{m}$
  $L_G \leftarrow \frac{1}{m}\sum_{i=1}^{m} -D_w(\tilde{\mathbf{x}}^{(i)}) + \lambda_2[I_{\text{target}}(\tilde{\mathbf{x}}^{(i)})\log O(\tilde{\mathbf{x}}^{(i)}) + (1 - I_{\text{target}}(\tilde{\mathbf{x}}^{(i)}))\log(1 - O(\tilde{\mathbf{x}}^{(i)})]$
  $\theta \leftarrow \text{Adam}(\nabla_\theta L_G, \theta, \alpha, \beta_1, \beta_2)$

#### 4.3 Architecture of GAN Models

For all the three GAN models (cWGAN, semi-supervised cWGAN, and semi-supervised gcWGAN) we use the same architecture for both the generator and the discriminator, as inspired by WGAN-GP (Gulrajani *et al.*, 2017). The generator and the discriminator's architectures in GAN are usually similar in computational expressiveness so that the training process could be stable, thus they are often transpose of each other. Both consist of residual blocks which include 2 layers of RELU for nonlinearity and 1D convolution with the skip connection between the input and the output.

As shown in the middle-right panel of Fig. S5, noise $\boldsymbol{z}$'s distribution as latent sequence space and fold representation $\boldsymbol{y}$ as the conditioning part are the inputs to the generator. Let $L$ be the maximal sequence length. The generator's architecture consists of the concatenation of $\boldsymbol{z}$ and $\boldsymbol{y}$ followed by a linear mapping to a higher-dimensional space ($L \times 500$). Five layers of residual blocks in sequence (20 layers in total) help improve the expressiveness of the model and their skip connections help avoid the gradient vanishing problem. Finally, 1D convolution brings down the dimensionality from $L \times 500$ to $L \times 21$ (20 characters for amino acids and one for padding); and a softmax layer produces a probability distribution over the characters for each position. The output of the generator is a continuous distribution over amino acids for each position. It can be converted to a discrete sequence by considering the character with the maximum probability for each position.

The generator's continuous distribution over the amino acids for each position with the fold representation are the inputs to the discriminator. The continuous distribution is used as the approximation for the one-hot encoding of discrete sequences because the latter, being non-differentiable, causes challenges in gradient calculation. First, a 1D convolutional layer converts the continuous distribution over amino acids ($L \times 21$) to a high dimension ($L \times 500$), followed by five layers of residual blocks to make the model deeper. Next, a linear layer brings the dimensionality down to 100. It is then concatenated with the fold representation and follow by the a fully connected layer with 300 neurons. Finally, a linear model converts the 300 dimensional space down to 1 dimension which is the output of the discriminator (or the critic in the WGAN).

### 5 Hyper-Parameter Tuning

#### 5.1 Criteria

##### 5.1.1 Convergence

The generator in cWGAN tries to minimize the loss function defined in Eq. 1 in the main text and the discriminator tries to maximize the loss. This min-max game originates from the zero-sum game in game theory. Since the generator seeks to minimize the loss, we can compare generators based on the total loss and the those with lower losses are more desirable. In WGAN, the total loss is named as “critic loss" since 1) the discriminator is named as the critic in WGAN and 2) the total loss is exactly the discriminator’s loss when we train the model (notice that the discriminator is present in all the three terms of $L_2$; however, the generator is only present in the second term).
To find out if the model has reached mathematical convergence, we assess the critic’s loss over hundreds of epochs.

##### 5.1.2 Nonsense Sequence Ratio

In our model, the length of the output of the generator or the input of the discriminator is fixed, but the length of the real sequences can be varied. So we fixed the output or input length to be the longest possible value (160 in the current study). And we used padding to fill the entries behind the sequence if its length is smaller than that value. One problem is that in the generated sequences, there can be padding appearing between or in front of the residue characters, which make it not resemble real sequences. We regarded these sequences as invalid or nonsense sequences, and called the ratio of the invalid ones in all generated sequences as the nonsense sequence ratio. Nonsense ratio was taken as one criteria to show the model performance in biological relevance. And for the generated sequences, firstly they need pass the sanity check; only those that pass the check are fed to the next-step and the nonsense sequences are discarded.

##### 5.1.3 Padding Ratio

A sequence that passes nonsense check above can still contain too much paddings to be relevant. We thus calculate the ratio of paddings over the whole sequence of characters and monitor them during model training.

##### 5.1.4 Sequence Novelty

Our final goal is to generate related sequences for some new folds in the future and also some new sequences for the known folds, do we would like the generated sequences are not exactly the training ones, which means we wanted the generated sequences to be novel. To measure the sequence novelty, we used the sequence identity as our criteria. The sequence identity is the ratio of the aligned characters in the whole alignment sequence, and we used the alignment score divided by the smallest alignment length to be the normalized alignment score so that it would be length independent. For each fold, we generated 100 valid sequences, and for each of the sequences, we calculated the sequence identity between it and all of representative sequence(s) related to the same fold, and then select the maximum ones. Then took the average of these selected scores on all the generated sequences of the fold as the final criteria.

#### 5.2 Hyper-parameters

##### 5.2.1 Initial learning rate

For the training process we used Adam Optimizer to optimize the loss function. Though for Adam Optimizer the learning is not fixed, the initial learning rate would influence it performance. This hyper-parameter is called learning rate in short for the following text. We set it to take values from 0.00001, 0.00005, 0.0001, 0.0002, to 0.0005, and fixed the critic iteration number and noise length to be 10 and 128 respectively. In figure S6 show that 1) although the learning rate of 0.0001 had the best critic loss and nonsense ratio, the padding ratio being 1 indicates all-padding sequences; 2) the learning rates 0.0001 and 0.0002 had similar performances and the former has a lower sequence identity. In the end, we choose 0.0001 as the learning rate and fix it for the next rounds of hyper-parameter tuning.

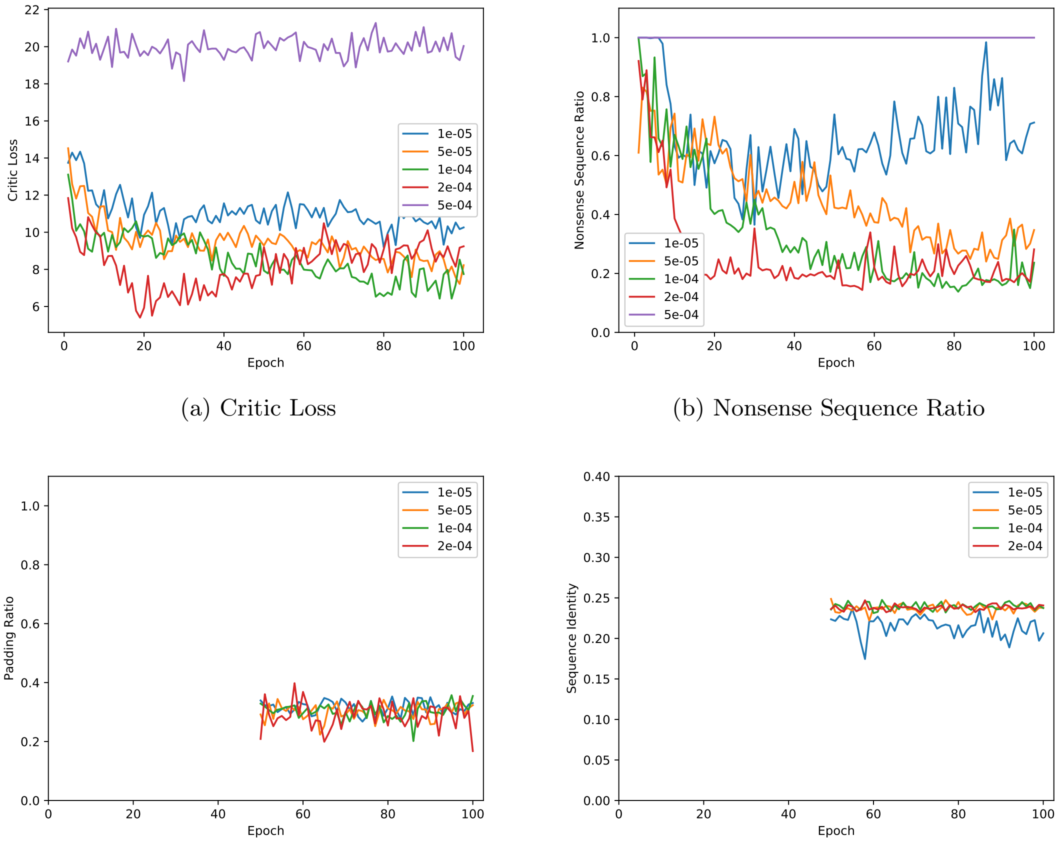


(a) Critic Loss

(b) Nonsense Sequence Ratio

(c) Padding Ratio

(d) Sequence Identity

Figure S6: Comparison for different initial learning rates when number of critic is fixed to 10 and noise length is fixed to 128

##### 5.2.2 Critic iteration number

For WGAN there is a generator and a critic, and we train the generator one time and the critic several times for each iteration. We fixed the initial learning rate to be 0.0001 and noise length to be 128, and then set the critic iteration number to be 5, 10 and 20 respectively. Based on Fig. S7, we found that when the number is 20 the model performed the best according to the critic loss and nonsense sequence ratio. Moreover, we can see that the sequence identity and padding ratio do not vary much for different critic iteration number. There, we selected 20 as the critic iteration number.

We use around 15 iterations per epoch and 1 batch (size of 64) for the generator. In each of the 15 iterations for the generator, the critic is updated 20 iterations with 20 batches. In total, we trained the whole gcWGAN model for 100 epochs.

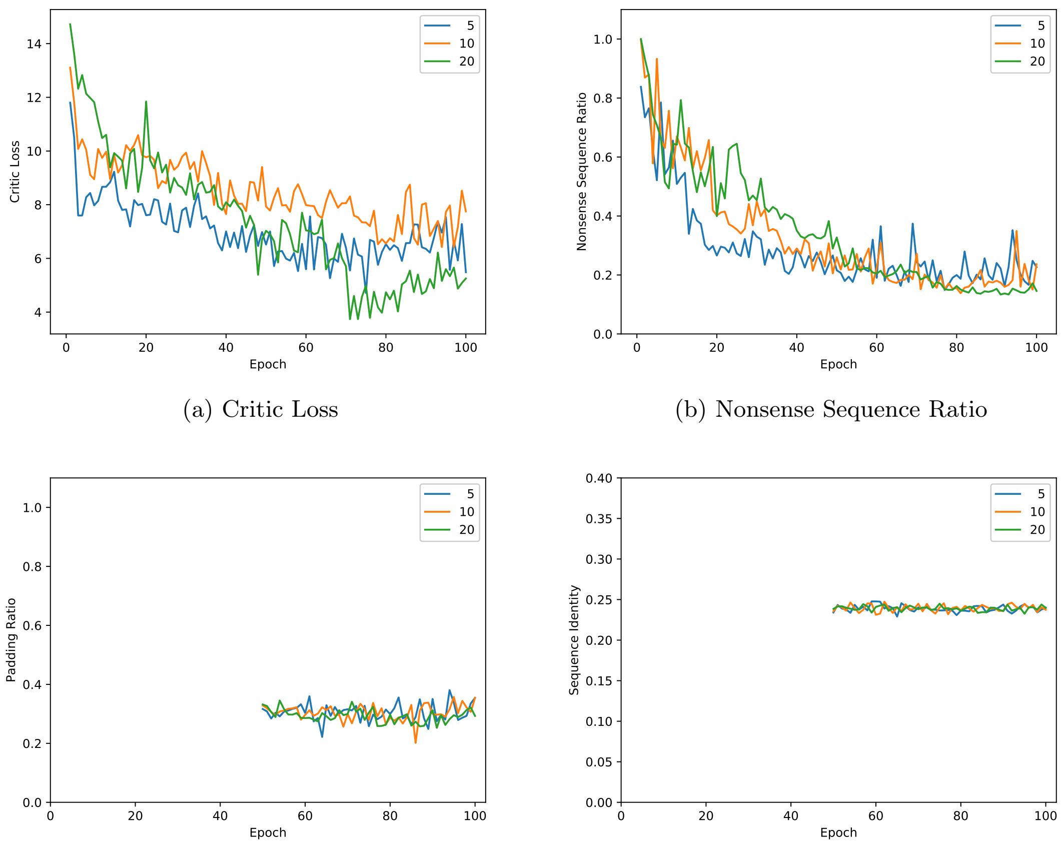


(a) Critic Loss

(b) Nonsense Sequence Ratio

(c) Padding Ratio

(d) Sequence Identity

Figure S7: Comparison for different initial learning rates when number of critic is fixed to 10 and noise length is fixed to 128

##### 5.2.3 Noise length

A random noise vector is an input to the generator and the generator maps the noise distribution to protein sequence distribution. The dimension of the vector is called noise length and also a hyper-parameter. We fixed the learning rate and critic iteration number to be the previous best ones and set the noise length to be 64, 128 and 256 in turn. Like the critic iteration number, we selected 128 as the final noise length according to the critic loss and nonsense sequence ratio based on Fig. S8. Moreover, we can see that the sequence identity and padding ratio do not vary much among the various lengths.

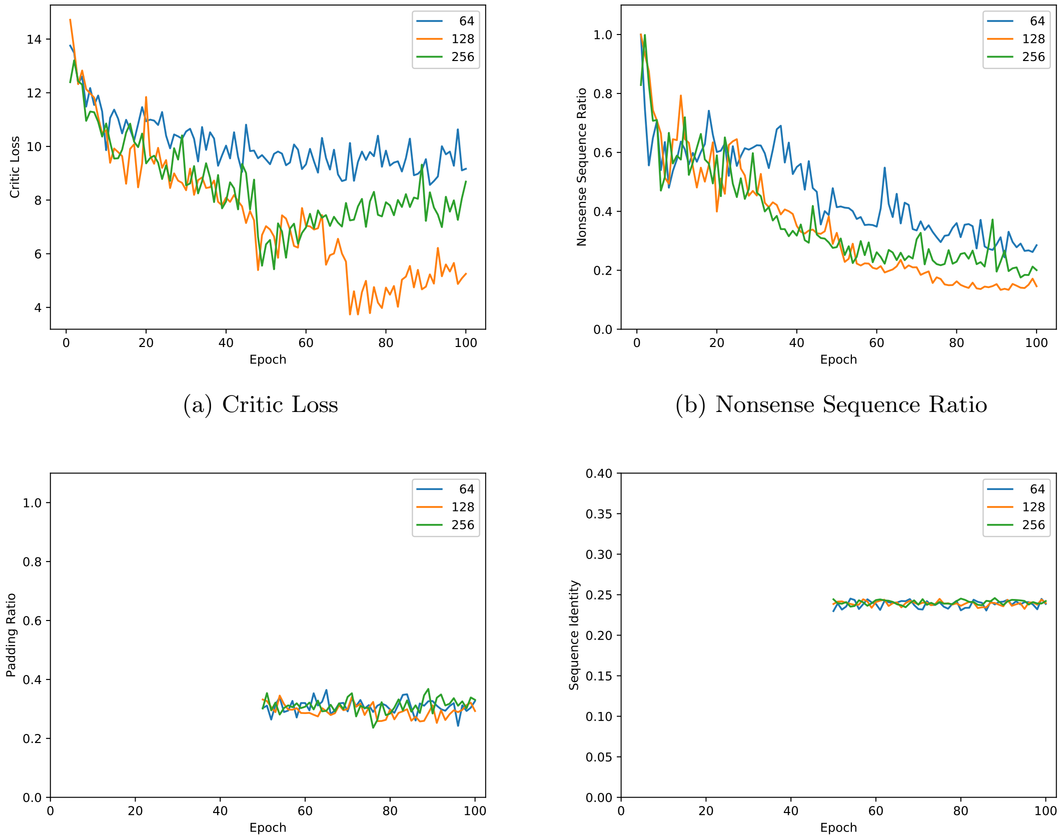


(a) Critic Loss

(b) Nonsense Sequence Ratio

(c) Padding Ratio

(d) Sequence Identity

Figure S8: Comparison for different noise length when number of critic is fixed to 10 and noise length is fixed to 128

##### 5.2.4 Feedback penalty in gcWGAN ($\lambda_2$)

For tuning the hyper-parameter ($\lambda_2$) corresponding to the feedback in gcWGAN, we consider the set of values from $\{0.001, 0.01, 0.1, 1, 10\}$. We observe that the model will diverge for values larger or equal than 0.1. We choose 0.01 for $\lambda_2$ based on the top-10 accuracy reported by the oracle for the validation folds. Specifically, for each validation fold, we generated 100 "valid" sequences using the generator trained at the last epoch and assessed their top-10 accuracy using the oracle.

### 6 Effect of Semi-supervision on gcWGAN

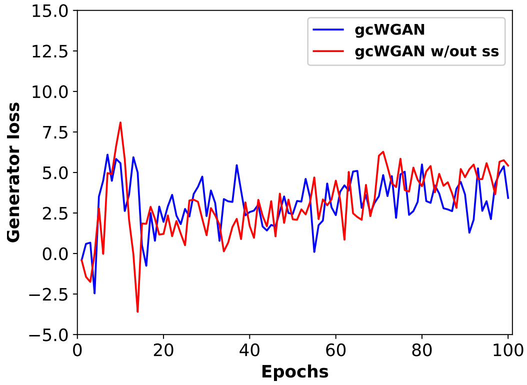


(a) Generator's Loss

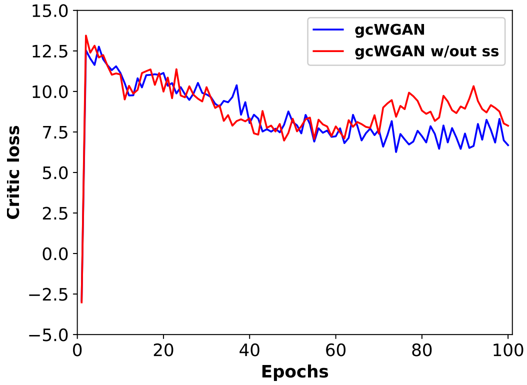


(b) Critic's Loss

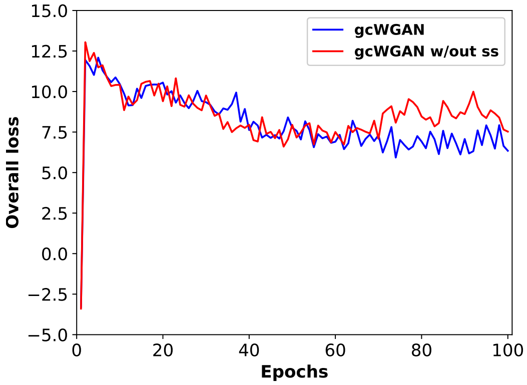


(c) Overall loss

Figure S9: Comparison of Guided-cWGAN training when initialized with semi-supervised cWGAN or cWGAN without semi-supervision (ss) for generator, critic and overall losses.

### 7 Assessing generated sequences prior to the final oracle check

Once the generator is trained and chosen, we first generated 100 "valid" sequences (passing the nonsense check) for each fold and examined their yield ratios (the accuracy according to the oracle's top-10 prediction). This is an intermediate sanity check as the final sequences will be only those that pass the oracle's check.

#### 7.1 gcWGAN with oracle feedback improves yields

| Yield Ratio | cWGAN | | | guided-cWGAN | | |
|---|---|---|---|---|---|---|
| | training | validation | test | training | validation | test |
| all | 1.3e-2 | 6.7e-4 | 2.1e-3 | 1.3e-2 | 9.2e-4 | 2.4e-3 |
| Fold Class Breakdowns | | | | | | |
| a | 2.0e-3 | 6.1e-5 | 1.2e-4 | 4.2e-3 | 1.9e-4 | 3.5e-4 |
| b | 2.8e-2 | 2.6e-3 | 7.0e-3 | 2.2e-2 | 2.0e-3 | 6.6e-3 |
| c | 4.6e-2 | 1.9e-3 | 9.8e-3 | 4.9e-2 | 3.2e-3 | 1.3e-2 |
| d | 5.2e-3 | 4.2e-4 | 7.8e-4 | 6.0e-3 | 9.5e-4 | 1.1e-3 |
| e | 1.1e-4 | $<$ 1e-5 | $<$ 1e-5 | 1.2e-3 | $<$ 1e-5 | $<$ 1e-5 |
| f | 4.9e-2 | 2.4e-4 | 4.1e-5 | 3.3e-2 | 4.4e-4 | $<$ 1e-5 |
| g | 2.1e-4 | 9.3e-6 | 2.5e-4 | 5.6e-4 | 1.4e-5 | 3.9e-4 |
| Sequence-Availability Class Breakdowns | | | | | | |
| easy | 3.5e-2 | 4.2e-3 | 2.1e-2 | 3.8e-2 | 7.9e-3 | 2.2e-2 |
| medium | 1.3e-2 | 2.1e-3 | 1.3e-3 | 1.1e-2 | 1.4e-3 | 1.4e-3 |
| hard | 2.0e-3 | 2.9e-4 | 3.6e-4 | 2.2e-3 | 4.2e-4 | 6.5e-4 |
| Oracle-Accuracy Class Breakdowns | | | | | | |
| $0 \sim 0.25$ | $<$ 1e-5 | 1.6e-5 | $<$ 1e-5 | 2.9e-5 | 1.1e-5 | $<$ 1e-5 |
| $0.25 \sim 0.5$ | 1.1e-3 | 2.2e-5 | $<$ 1e-5 | 7.8e-4 | 5.1e-5 | 2.6e-5 |
| $0.5 \sim 0.75$ | 2.5e-3 | 1.6e-3 | 1.3e-3 | 4.3e-3 | 3.1e-3 | 2.1e-3 |
| $0.75 \sim 1$ | 5.1e-2 | 3.2e-3 | 1.7e-2 | 4.8e-2 | 4.6e-3 | 1.9e-2 |

Table S5: Yield ratio results for cWGAN and guided cWGAN

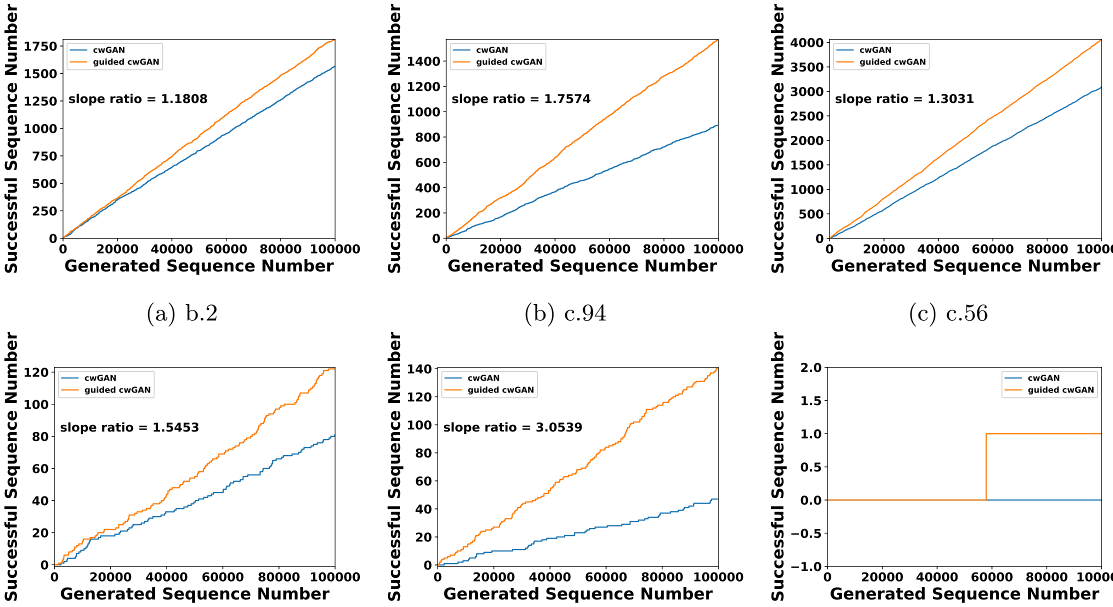


(a) b.2 (b) c.94 (c) c.56

(d) a.35 (e) g.44 (f) d.107

Figure S10: The number of “yields" versus the number of sequences generated by cWGAN/gcWGAN for selected folds.

#### 7.2 cWGAN and gcWGAN have higher yields for more folds than cVAE

| | cVAE | | cWGAN | | gcWGAN | |
|---|---|---|---|---|---|---|
| | yield ratio | fold coverage | yield ratio | fold coverage | yield ratio | fold coverage |
| all | 7.4e-3 | 0.075 | 2.1e-3 | 0.318 | 2.4e-3 | 0.345 |
| Fold Class Breakdowns | | | | | | |
| a | $<$ 1e-5 | 0 | 1.2e-4 | 0.286 | 3.5e-4 | 0.357 |
| b | 5.2e-2 | 0.200 | 7.0e-3 | 0.333 | 6.6e-3 | 0.333 |
| c | 1.6e-4 | 0.375 | 9.8e-3 | 0.750 | 1.3e-2 | 0.750 |
| d | $<$ 1e-5 | 0.024 | 7.8e-4 | 0.268 | 1.1e-3 | 0.317 |
| e | $<$ 1e-5 | 0 | $<$ 1e-5 | 0 | $<$ 1e-5 | 0 |
| f | $<$ 1e-5 | 0 | 4.1e-5 | 0.667 | $<$ 1e-5 | 0 |
| g | $<$ 1e-5 | 0.091 | 2.5e-4 | 0.182 | 3.9e-4 | 0.273 |
| Sequence-Availability Class Breakdowns | | | | | | |
| easy | 5.4e-4 | 0.500 | 2.1e-2 | 1 | 2.2e-2 | 0.875 |
| medium | 3.5e-2 | 0.136 | 1.3e-3 | 0.500 | 1.4e-3 | 0.455 |
| hard | 1.4e-5 | 0.013 | 3.6e-4 | 0.195 | 6.5e-4 | 0.260 |

Table S6: Yield comparison among cVAE, cWGAN and guided-cWGAN for the test set.

| Cases | cVAE | cWGAN | gcWGAN |
|---|---|---|---|
| b.2 | 2.2e-4 | 1.9e-2 | 1.6e-2 |
| c.94 | 3.0e-4 | 1.3e-2 | 1.4e-2 |
| c.56 | 1.1e-3 | 2.5e-2 | 4.2e-2 |
| a.35 | $<$ 1e-5 | 6.6e-4 | 1.5e-3 |
| g.44 | 2.0e-5 | 1.5e-3 | 1.3e-3 |
| d.107 | $<$ 1e-5 | 6.5e-5 | 4.4e-5 |

Table S7: Yield ratio for 6 selected representative folds for cVAE, cWGAN and guided-cWGAN

### 8 Assessing final sequence designs

For each of the six representative folds and a novel fold, we generated 100 valid sequences that pass the final check of the oracle (modified DeepSF). For each fold, we "folded" the first ten sequences, predicted 100 structures for each fold using Rosetta, and evaluated their structural accuracy compared to the known representative structure using TM score. We also evaluated sequence diversity among each set of 100 sequences and sequence novelty of them compared to the known representative sequence.

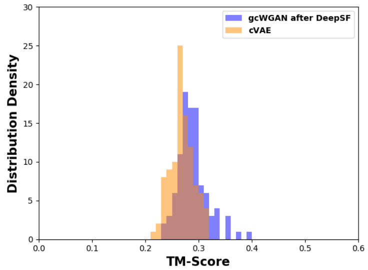


(a) b.2

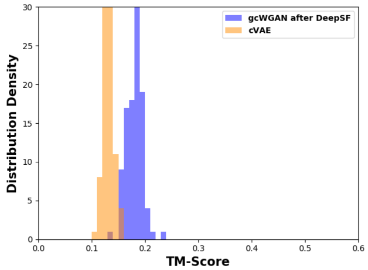


(b) c.94

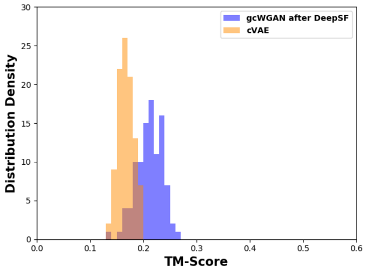


(c) c.56

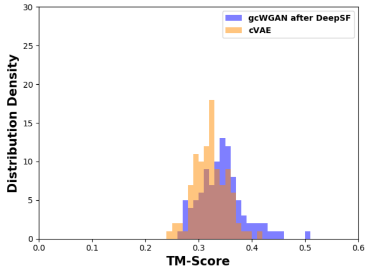


(d) a.35

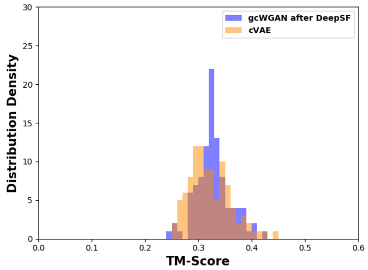


(e) g.44

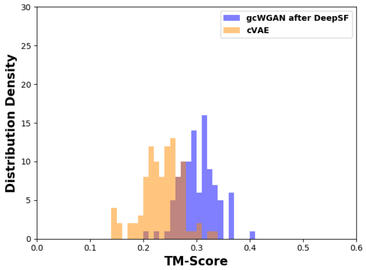


(f) d.107

Figure S11: Distribution of the TM scores between the selected predictions and the ground truth on the 6 selected cases.

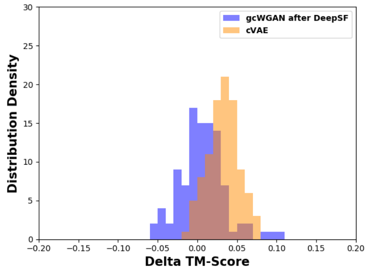


(a) b.2

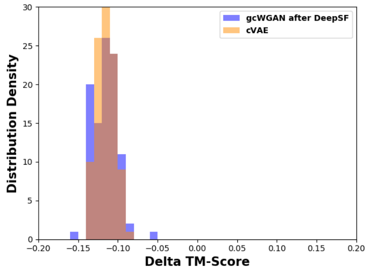


(b) c.94

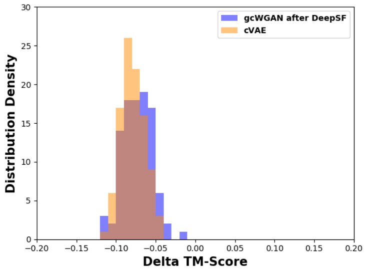


(c) c.56

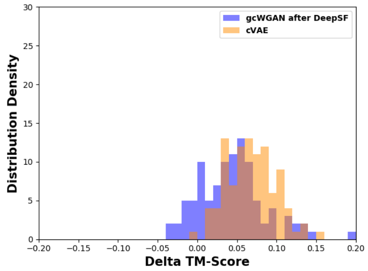


(d) a.35

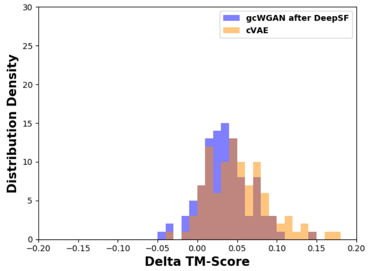


(e) g.44

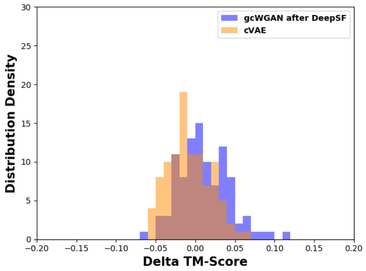


(f) d.107

Figure S12: Distribution of ΔTM scores between the selected predictions and the ground truth on the 6 selected cases.

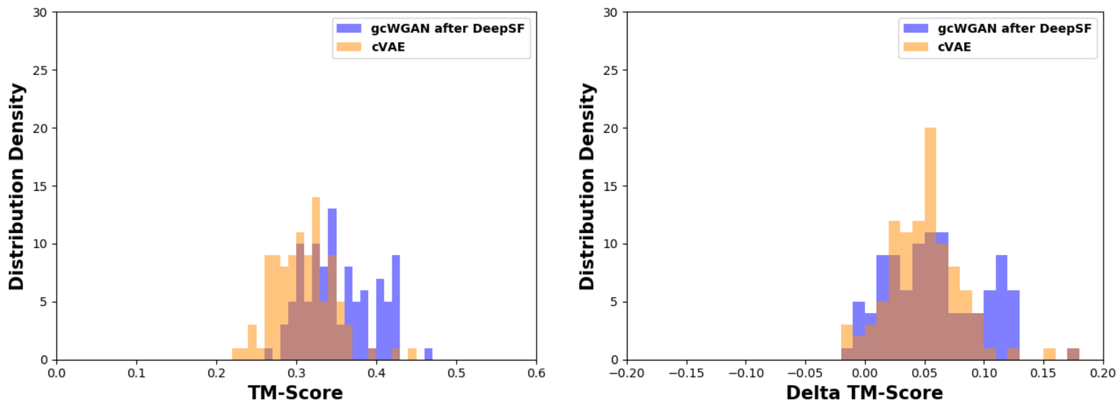


(a) TM score

(b) Delta TM score

Figure S13: Distribution of TM scores and delta TM scores between the selected predictions and the ground truth on the novel fold.

| | TM scores | | delta TM scores | |
|---|---|---|---|---|
| | T value | p value | T value | p value |
| novel fold | 7.795 | 1.923e-13 | 1.825 | 3.477e-2 |
| b.2 | 5.548 | 4.943e-8 | -7.339 | 1.000 |
| c.94 | 26.758 | 5.186e-62 | 2.846e-2 | 4.887e-1 |
| c.56 | 15.474 | 5.477e-34 | 2.339 | 1.018e-2 |
| a.35 | 3.938 | 5.899e-5 | -4.713 | 1.000 |
| g.44 | 1.398 | 8.182e-2 | -3.474 | 9.997e-1 |
| d.107 | 13.391 | 9.438e-30 | 4.735 | 2.124e-6 |

Table S8: One-sided $t$-test results for the distributions in TM-scores and $\Delta$TM-scores .

#### 8.1 Structure alignment between the ground truth and the best prediction of gcWGAN designs

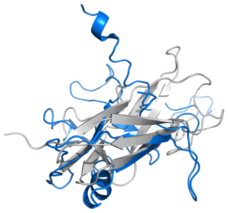

(a) b.2 (TM score = 0.3920)

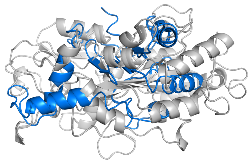

(b) c.94 (TM score = 0.2378)

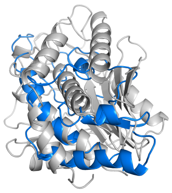

(c) c.56 (TM score = 0.2682)

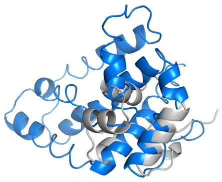

(d) a.35 (TM score = 0.5060)

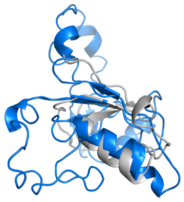

(e) g.44 (TM score = 0.4205)

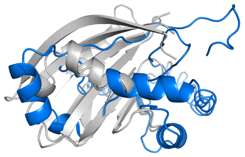

(f) d.107 (TM score = 0.4095)

Figure S14: Protein alignment for the 6 selected cases.

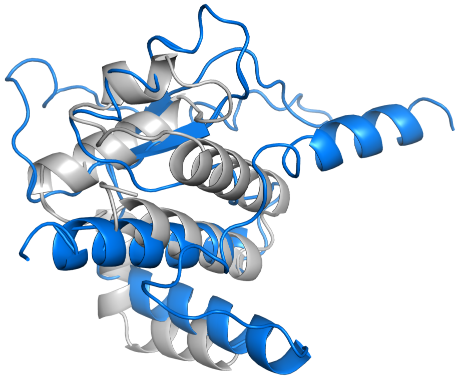

Figure S15: Protein alignment for the novel fold (TM score = 0.4700).

### 9 Diversity of sequences generated by cWGAN and cVAE

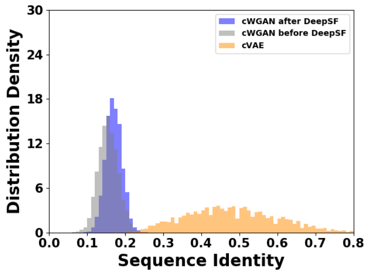


(a) b.2

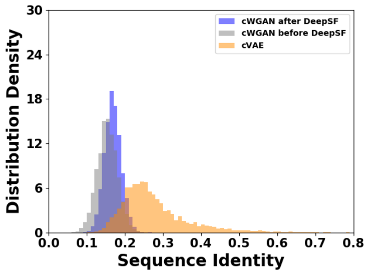


(b) c.94

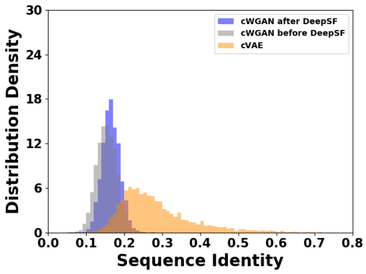


(c) c.56

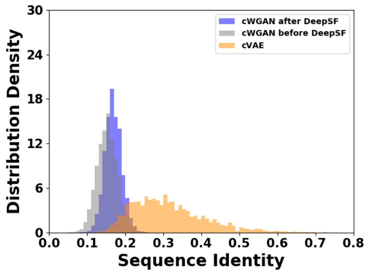


(d) a.35

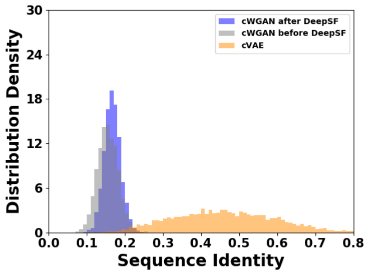


(e) g.44

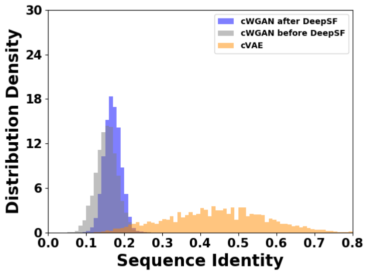


(f) d.107

Figure S16: Sequence diversity for the 6 selected cases based on cWGAN

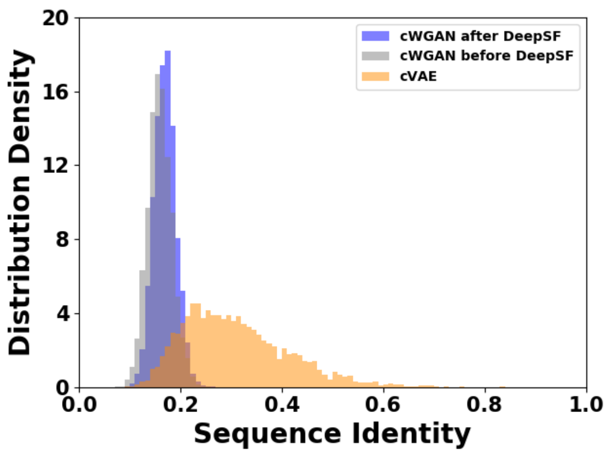


Figure S17: Sequence diversity for the novel fold based on cWGAN

### 10 Novelty of sequences generated by cWGAN and cVAE

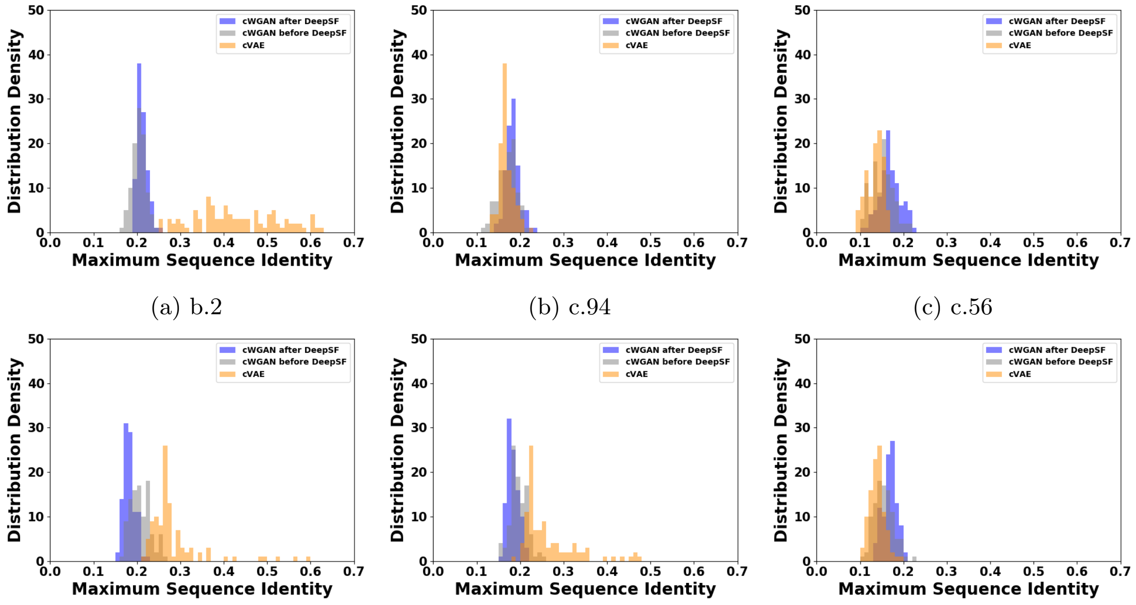


(a) b.2 (b) c.94 (c) c.56

(d) a.35 (e) g.44 (f) d.107

Figure S18: Sequence novelty distribution for the 6 selected cases based on cWGAN

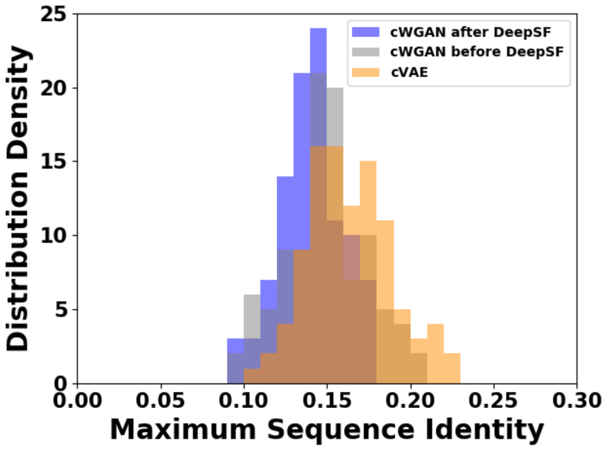


Figure S19: Sequence novelty distribution for the novel fold based on cWGAN
